# RICOPILI: Rapid Imputation for COnsortias PIpeLIne

**DOI:** 10.1101/587196

**Authors:** Max Lam, Swapnil Awasthi, Hunna J. Watson, Jackie Goldstein, Georgia Panagiotaropoulou, Vassily Trubetskoy, Robert Karlsson, Oleksander Frei, Chun-Chieh Fan, Ward De Witte, Nina R. Mota, Niamh Mullins, Nora Skarabis, Hailiang Huang, Benjamin Neale, Mark Daly, Manuel Mattheissen, Raymond Walters, Stephan Ripke

**Author notes:** These authors contributed equally (Max Lam, Swapnil Awasthi).

## Abstract

**Motivation:** Genome-wide association study (GWAS) analyses, at sufficient sample sizes and power, have successfully revealed biological insights for several complex traits. RICOPILI, an open sourced Perl-based pipeline was developed to address the challenges of rapidly processing large scale multi-cohort GWAS studies including quality control, imputation and downstream analyses. The pipeline is computationally efficient with portability to a wide range of high-performance computing (HPC) environments.

**Summary:** RICOPILI was created as the Psychiatric Genomics Consortium (PGC) pipeline for GWAS and has been adopted by other users. The pipeline features i) technical and genomic quality control in case-control and trio cohorts ii) genome-wide phasing and imputation iv) association analysis v) meta-analysis vi) polygenic risk scoring and vii) replication analysis. Notably, a major differentiator from other GWAS pipelines, RICOPILI leverages on automated parallelization and cluster job management approaches for rapid production of imputed genome-wide data. A comprehensive meta-analysis of simulated GWAS data has been incorporated demonstrating each step of the pipeline. This includes all of the associated visualization plots, to allow ease of data interpretation and manuscript preparation. Simulated GWAS datasets are also packaged with the pipeline for user training tutorials and developer work.

**Availability and Implementation:** RICOPILI has a flexible architecture to allow for ongoing development and incorporation of newer available algorithms and is adaptable to various HPC environments (QSUB, BSUB, SLURM and others). Specific links for genomic resources are either directly provided in this paper or via tutorials and external links. The central location hosting scripts and tutorials is found at this URL: https://sites.google.com/a/broadinstitute.org/RICOPILI/home

**Supplementary information:** Supplementary data are available.

## 1. Introduction

Genome-wide association studies (GWASs) have enabled the discovery of genetic variants underlying a plethora of complex traits^1^. GWASs have highlighted previously unknown biological mechanisms associated with complex diseases and traits (Breen *et al.*, 2016). The Psychiatric Genomics Consortium (PGC) (http://www.med.unc.edu/pgc) the largest umbrella organization for psychiatric genetics (Sullivan *et al.*, 2017) - have made possible to advance the objectives of a) revealing biological insights of psychiatric illness, b) informing clinical practice, and c) presenting new therapeutic targets through sheer number of cohorts for GWASs across various psychiatric traits (Breen *et al.*, 2016; Sullivan *et al.*, 2012). The exponential availability of cohorts, requires efficient, consistent and standardized approaches for various aspects of GWAS data management and analysis. Here, we introduce RICOPILI, the pipeline that automates rapid GWAS analysis workflow across various PGC workgroups. The pipeline is state-of-art, constantly incorporating latest available GWAS computational techniques and methods. With open-sourced simulated GWAS datasets and training tutorials packaged with the pipeline, RICOPILI is ideal for those contributing to large-scale genetic studies. To our understanding, RICOPILI is the only GWAS pipeline allowing secure data management, efficient data processing, and downstream analysis scalable on both desktop and cluster environments (Supplementary Table 1)

## 2. Design and Implementation

### 2.1 Pipeline description

RICOPILI automates and integrates standard GWAS analysis methods, allowing for automated cluster submission and parallelization. The pipeline unifies standard software for its functions and implements best data analysis practices, provides sensible default settings while permitting the user to flexibly customize filters, thresholds, and job resources as required. The optimization of cluster resources allows computations and visualizations to be completed quickly without significant user intervention. Written prediminantly in Perl and R, the pipeline is organized according to analysis modules. Each module runs in its entirety via a single command line. The main module functions include:

- Pre-imputation technical quality control (QC);
- Principal components analysis (PCA) and relatedness estimation;
- Genome-wide imputation of genotype probabilities and generation of best guess genotypes;
- Downstream analyses, including GWAS, and meta-analysis, and polygenic risk scoring;
- Harmonizing large imputation reference panels (such as 1000 Genomes, and the Haplotype Reference Consortium) to fit the architecture of RICOPILI.

RICOPILI takes data set with unfiltered genotype calls, through trait association analysis, multi-cohort meta-analysis, linkage disequilibrium (LD) score regression (Bulik-Sullivan *et al.*, 2015), conditional analysis, replication analysis, and polygenic risk scoring (Supplementary Figure 1). Little intermediate interaction is required, allowing for efficient standardized analysis of genome-wide data and results. Standardized file naming conventions are designed to optimize overview and analysis record tracking within large-scale genetic projects. Publication-ready data visualizations and reports (in PDF and Excel format) permits easy evaluation of the results. Simulated datasets are also available with the pipeline for training and development purposes. In the ensuing sections, we describe the main components of the pipeline.

### 2.2 Pre-imputation/quality control (Supplementary Figure 2)

The pre-imputation/QC module (Supplementary Section 1) consists of the following, general steps:

- Inferring the genotyping chip;
- Standardizing file names and sample identifiers, incorporating chip information, and ensuring that sample IDs across distinct cohorts are unique while keeping original sample IDs intact;
- Carrying out technical sample and variant QC procedures.

Detailed sample and variant filtering reports provide diagnostics to identify possible QC issues and solutions. Quality controlled datasets are saved separately for downstream analysis.

### 2.3 Principal components analysis (Supplementary Figure 3)

- The PCA module (Supplementary Section 2) fulfills two objectives:
- Identify and remove duplicated or related samples for case-control and trio cohorts;
- Assess ancestral outliers and population stratification with EIGENSTRAT (Price *et al.* 2006);
- Principal component scores are computed, and could be utilized for visualization or as covariates to adjust for population structure in downstream post-imputation GWAS.

### 2.4 Imputation (Supplementary Figure 4)

RICOPILI automates computationally costly genotype imputation with an optimized routine for HPC environments (Supplementary Section 3). This module aligns genotype data to the imputation reference, pre-phases haplotypes, and executes imputation. Users have the option to:

- Impute genotypes to the 1000 Genomes (1000 Genomes Project Consortium *et al.* 2015) or Haplotype Reference Consortium panel (McCarthy *et al.*, 2016);
- Perform pre-phasing with Eagle (Loh *et al.*, 2016) or SHAPEIT (Delaneau *et al.*, 2012);
- Perform imputation with IMPUTE (Bycroft *et al.*, 2018; Howie *et al.*, 2009) or Minimac (Das *et al.*, 2016; Howie *et al.*, 2012).

RICOPILI allows for automated data preparation, alignment and sharing with public imputation servers^2^ (e.g. Michigan^3^, Sanger^4^), and reintegration of the results back into the RICIOPILI data structure. This is especially beneficial if an HPC environment is not accessible, and imputation by third party services has been approved by the user’s local Institutional Review Board (IRB). The outputs are a set of genotype probabilities for all markers and ready-for-analysis “best-guess” genotype hardcall files filtered on imputation quality and minor allele frequency. RICOPILI allows the creation of case-pseudo-controls to handle imputation and association procedures for trios.

### 2.5 Post-imputation (Supplementary Figure 5)

The post-imputation module (Supplementary Section 4) performs association analysis using imputed dosage files, meta-analysis via METAL (Willer *et al.*, 2010), conditional analysis, polygenic risk scoring, LD score regression (Bulik-Sullivan *et al.*, 2015) and replication analysis. Covariates (e.g., age, sex, principal components from PCA) and alternative phenotypes, including quantitative traits may be incorporated within the post-imputation module. Automated “clumping” of genome-wide significant SNPs to facilitate identification of independently associated genetic loci. Publication-ready reports and visualizations such as Manhattan plots, QQ-plots, forest plots, annotated region plots, and polygenic risk distributions are generated by the module as well.

### 2.6 Additional utility modules

RICOPILI allows for additional features and modules (see Supplementary Information). Including, (1) reference builder: builds reference data for genotype imputation from publicly accessible reference panels (Supplementary Figure 6) (2) replication of GWAS: using external summary data or those generated by RICOPILI (3) polygenic leave-one-out analysis: where each input data set is used as a hold out and polygenic risk prediction is done iteratively across hold out data.

### 2.7 Availability of simulated GWAS data (Supplementary Section 6)

To allow new users to familiarize themselves with RICOPILI and experienced users to develop new functionality for the pipeline, we simulated freely available GWAS data using HAPGEN (Su *et al.*, 2011). The data set comprises 6,200 “individuals” across ~600,000 markers based on the Illumina OmniExpress, a widely used genotyping platform. For training and development purposes, population stratification, cross-sample relatedness, and technical errors were introduced to the simulated data. The sample is separated into five datasets “HapGen5” packaged with RICOPILI^5^. Data description and results are described in further detail in <u>Extended Data Analysis and User Guide</u>.

### 2.8 Cluster portability (Supplementary Section 7)

RICOPILI is portable^6^ to various LINUX-based HPC environments (e.g. BSUB^7^, QSUB^8^, SLURM, GCP (Google Cloud Platform^9^)). In the absence of a HPC environment, RICOPILI can use the full potential of multi-core machines with parallel optimization. Regular updates and maintenance of the pipeline are carried out to incorporate the latest advances in genetic association methods. Ongoing support includes an active user forum (https://groups.google.com/forum/#!forum/ricopili-user-group), support website (https://sites.google.com/a/broadinstitute.org/ricopili/home), and detailed tutorials written by current RICOPILI analysts (consult footnotes).

## 3. Discussion

RICOPILI has supported the analytical capability of the PGC, encompassing over 800 investigators internationally. The consortium is a testament to collaborative science that has unified much of the field and collated data collections, and enabled rapid progress in uncovering the genetic and biological basis of psychiatric disorders. RICOPILI addresses the need for a rapid computational pipeline for GWAS that integrates leading bioinformatics resources and produces publication-ready outputs. The PGC has reported GWAS studies in high-impact publications, most of which featured RICOPILI as the main analysis pipeline – including the seminal report identifying 108 GWAS loci for schizophrenia (Ripke *et al.*, 2014). The pipeline has been adapted across various consortia, with 112 analysts performing rapid computation for GWAS to date. For this reason, we introduce RICOPILI to an audience of principal investigators, academics, analysts, and all personnel tasked with determining the common variation underlying complex, heritable diseases and traits.

## Acknowledgments

Computing and network infrastructure was provided by various sources, including SURFsara (Genetic Cluster Computer; LISA cluster) and the Stanley Center for Psychiatric Research at the Broad Institute of MIT and Harvard.

## Footnotes

1 https://www.ebi.ac.uk/gwas/diagram

2 https://docs.google.com/document/d/18dupvU4kw11slREc1TUfwQwhO_eI0n_MeKVpwi4HLNA/

3 https://imputationserver.sph.umich.edu/index.html#!pages/home

4 https://imputation.sanger.ac.uk/

5 https://docs.google.com/document/d/1ux_FbwnvSzaiBVEwgS7eWJoYlnc_o0YHFb07SPQsYjI/

6 https://docs.google.com/document/d/14aa-oeT5hF541I8hHsDAL_42oyvlHRC5FWR7gir4xco/

7 https://docs.google.com/document/d/1fNFnC3-rBZkmtH47Je_yUfGatB9qhDGi9HtMSA3_MPw/

8 https://docs.google.com/document/d/1oY5IA4a6yG_pmbvWJC8A6MTzjYoGzVlqQ_aXUwWCl8I/

9 https://cloud.google.com/

