## Supplementary for "RICOPILI: Rapid Imputation for COnsortias PIpeLIne"

### Supplementary Information

**Title:** RICOPILI: **R**apid **I**mputation for **CO**nsortias **PI**pe**LI**ne

**Supplementary Sections**

Further detailed descriptions of the pipeline, in-depth discussion of analysis steps, and frequently asked questions (FAQs) are available on the RICOPILI website[1]. The RICOPILI PIPELINE and accompanying website are maintained regularly and updated to reflect the latest advancements in genome-wide association study (GWAS) methodology. A RICOPILI users' group forum is available at https://sites.google.com/a/broadinstitute.org/ricopili/users-section/. The sections of this supplement are:

1. Pre-Imputation/Quality Control (QC) Module Description
2. Principal Components Analysis Module Description
3. Imputation Module Description
4. Post-Imputation Module Description
5. Additional Utility Modules
6. HAPGEN Data Simulation Methods and Detailed Tutorial to Perform Quality Control, Imputation, Association and Meta-analyses
7. Cluster Portability
8. RICOPILI Output Logging Stages

**Supplementary Table**

Supplementary Table 1. Pipeline Comparisons with RICOPILI

**Supplementary Figures**

Supplementary Figure 1. RICOPILI visualizations
Supplementary Figure 2. Pre-Imputation/Quality Control module workflow
Supplementary Figure 3. Principal Components Analysis (PCA) module workflow
Supplementary Figure 4. imputation module workflow
Supplementary Figure 5. Post-Imputation module workflow
Supplementary Figure 6. Reference builder module workflow

[1] https://sites.google.com/a/broadinstitute.org/ricopili/home (RICOPILI Home Page)

#### 1. Pre-Imputation/Quality Control (QC) Module Description

The pipeline begins with quality control (QC) of raw datasets in PLINK binary format (Purcell *et al.*, 2007). Disease-group, population- and case-control-assignment and platform information is appended to the family identifier (FID) to create cross-cohort uniqueness and integrate basic meta-information. Next, technical QC is applied. A pre-filter 95% variant call rate is applied to avoid excessive sample loss in the next step, especially after large scale data merges of cohorts from different sources. This is followed by sample filtering to retain high-quality samples, with default filters and thresholds applied that can be adjusted flexibly by the user. The defaults include a genotyping call rate greater than 98% and inbreeding coefficient (F) less than +-0.2. Sex checks on the X-chromosome provoke exclusions (F_male < 0.5, F_female > 0.5) and warnings (F_male < 0.8, F_female > 0.2). This step is especially crucial in identifying problematic handling of specimens during the genotyping process. For trio data, individuals with more than 10,000 Mendelian errors are excluded. Subsequent variant-level filtering is performed to retain high-quality variants, again with default filters and thresholds that can be adjusted by the user. Default filtering excludes variants with more than 2% missing-rate, variants with differences in call rates between cases and controls greater than 2%, monomorphic variants, and Hardy-Weinberg Equilibrium (HWE) deviations with a significance less than $P < 10^{-10}$ (cases) and $P < 10^{-6}$ (controls). An initial association analysis without covariates is performed to identify genome-wide inflation and other indicators of remaining QC artifacts. Comprehensive diagnostic plots and reports summarize sample and variant exclusion outcomes. The researcher can use this information to assess the quality of the raw and quality-controlled data and evaluate additional remediation that may be required. The tutorial section with simulated data shows many commonly observed technical problems and also presents their solutions in RICOPILI[2].

#### 2. Principal Components Analysis Module Description

The PCA module workflow begins with extracting a subset of high-quality, high-frequency autosomal SNPs to use for relatedness and population testing. First PLINK binary files from all cohorts to be included in the PCA are aligned and merged and linkage disequilibrium (LD) pruned. Long-range LD regions - specifically the major histocompatibility complex (MHC) in chromosome 6 (chr 6, 25-35 MB) and the inversion in chromosome 8 (chr 8, 7-13 MB) - are removed to avoid artificial PCA creation from single genotypes in these genomic areas. SNPs at HWE thresholds at $P < 10^{-3}$ and markers with minor allele frequency (MAF) < 5% are also excluded. This results typically in a final set of 3,000 to 100,000 SNPs, depending on the variety of different genotyping platforms used. The PCA module within RICOPILI fulfills dual roles. First, the PCA module may be applied to single cohorts along with the pre-imputation/QC module to adjust for lambda inflation due to population outliers confounded with phenotype associations. Second, the outputs from the PCA module are able to be used in the later stages of association analysis to build covariates for adjusting for fine-grained population stratification, and cross-cohort duplication and relatedness. The PCA module automatically removes duplicate samples for downstream analysis. Caution is however recommended in PCA analysis of greater than 40 cohorts which may result only in a low number of overlapping SNPs across cohorts due to differing genotyping platforms. In such extreme cases of low SNPs emerging from PCA, we recommend the approach of using i) de-duplicated ID definition from cross-cohort combined PCA, and ii) applying single cohort principal components as covariates.

[2] https://docs.google.com/document/d/1ux_FbwnvSzaiBVEwgS7eWJoYlnc_o0YHFb07SPQsYjI/ (RICOPILI Tutorial)

##### *2.1. Calculation of Sample Relatedness*

Relatedness indices between all pairs of individuals (intra- and cross-cohort) are generated from the above SNP set within PLINK. To facilitate relatedness testing of large sample sizes, the combined dataset is split into chunks of 200 individuals. With a sufficiently large number of SNPs (~100,000 independent SNPs) it is possible to estimate the pairwise identity-by-descent (IBD) for each pair of individuals within the cohort. Across these SNPs, it is possible to generate based on allele frequencies, the probability of individuals having two identical homozygous alleles or sharing a single heterozygous allele. The proportion of IBD or PiHat could be defined as the sum of the probability of individuals sharing two homozygous alleles and half of the probability of sharing a heterozygous allele (Purcell *et al.*, 2007). Typically, identical twins or duplicated samples share 100% of IBD (PiHat = 1), first degree relatives share 50% (PiHat = 0.5), second degree relatives share 25%, and third degree relatives share 12.5%. Only sample pairs with PiHat > 0.1 are reported. All pairs of individuals with PiHat > 0.2 are identified and one each is removed from the PCA step. Trio samples are prioritized for retention, followed by cases, followed by samples genotyped on a preferred platform. During the process, already excluded individuals are prioritized for exclusion. Otherwise the exclusion decision is random. As a result, a collection of unrelated individuals with optimized cohort size is carried forward to the calculation of ancestral principal components.

##### *2.2 Calculation of Ancestral Components*

The ancestral multidimensional scaling procedures conduct a series of variant allele alignment before conducting SmartPCA in fastmode (Price *et al.*, 2006; Galinsky, Bhatia, *et al.*, 2016; Galinsky, Loh, *et al.*, 2016). The procedure generates 20 principal components. Several PCA plots are produced, including single- and two-dimensional plots, which visually differentiate cases and controls by color, to allow for comprehensive data visualization. These can be used to visually identify outlying individuals and evaluate whether ancestry is balanced among cases and controls. RICOPILI includes the option to overplot a public reference dataset (i.e., from the 1000 Genomes Project) to facilitate characterizing ancestry of within the analyzed sample.

The PCA covariates file generated at this point within the module is typically used in downstream GWAS containing unrelated individuals. The principal components from formerly excluded individuals (during the sample relatedness process above) are inherited from their included partners. Hence, the user can decide to use this file in combination with a different sample-inclusion/exclusion definition. The user would be able to identify and remove ancestral outliers, repeat PCA and the QC module to observe the effect of such action on the association analysis and QC matrices.

### 3. Imputation Module Description

RICOPILI automates genotype imputation, giving users options for various reference panels, being successfully tested with HapMap2 (International HapMap Consortium, 2007), HapMap3[3] (International HapMap 3 Consortium, 2010), human leukocyte antigens (HLA) imputation reference[4] (Jia *et al.*, 2013), 1000 Genomes Phase 1 (1000 Genomes Project Consortium, 2010) and Phase 3[5] (1000 Genomes Project Consortium, 2015), and the Haplotype Reference Consortium (HRC) panel (McCarthy *et al.* 2016) available through the European Genome-phenome Archive (EGA) (accession number EGAS00001001710). Users are given the option of performing imputation per cohort, or imputing across multiple cohorts together.

The imputation module first converts the input data to hg19 (Human Build GRCh37) using Liftover (Hinrichs *et al.*, 2006) if the pipeline detects data from a different build. SNP name inconsistencies are resolved, alleles are aligned, strand flips are adjusted for strand unambiguous variants as well as for strand ambiguous SNPs, the latter via comparing allele frequencies with the selected reference ancestry, whilst removing very common (default: MAF > 40%) ambiguous SNPs. All variants with excessive frequency difference (default: >15%) to the selected reference ancestry are removed.

For optimal computational efficiency, pre-phasing haplotypes from the genotypes and subsequent imputation is done in genomic chunks of variable sizes (defined during the reference building process). Be default, the public HRC reference is divided into 132 genomic chunks with around 287,000 variants each. The user can choose from a variety of different chunks sizes and hence choose for higher or lower parallelization.

Pre-phasing is performed with Eagle (Loh *et al.*, 2016), and imputation with Minimac3 (Das *et al.*, 2016). Optionally, the user can also choose the SHAPEIT/IMPUTE phasing/imputation combination (Bycroft *et* al., 2018; Delaneau *et al.*, 2012; Howie *et al.*, 2009). The latter is not as computationally efficient but can provide a better handling of trio phasing and imputation[6,7].

After very light quality filtering (defaults: imputation information score (INFO) > 0.1, MAF > 0.005), the output is a set of genotype probabilities for all markers. In addition, ready-for-analysis "best-guess" genotype files are created bycalling the genotype with a probability of > 0.8 (if no genotype probability reaches that threshold, the call is set to missing). Best guess genotypes are provided in three forms: (1) no additional filter, (2) missing rate < 2%, (3) missing rate <1% and MAF >5%).

RICOPILI allows the creation of case-pseudo controls to handle the imputation and association procedures for trios. After phasing the trios the pseudo-controls are built from the non-transmitted alleles. From that point on, imputation of the affected offspring and each perfectly matched pseudo-control is pursued in the above described process.

---

[3] https://www.ncbi.nlm.nih.gov/variation/news/NCBI_retiring_HapMap/
[4] http://software.broadinstitute.org/mpg/snp2hla/
[5] http://www.internationalgenome.org/data
[6] https://sites.google.com/a/broadinstitute.org/ricopili/imputation (RICOPILI Imputation)
[7] https://docs.google.com/document/d/1juiqszi-9Wg7vei0_d7QHT7G_a5QHQUbxsoYPVLMyTo/ (Trio Imputation)

In recent versions of RICOPILI (Jan 29th 2019 and later), the imputation module is integrated with publicly available genotype imputation services e.g., Michigan Imputation Server[8] or Sanger Imputation Service[9]. Using the --deploy function, users without the benefit of cluster access may first perform pre-imputation and PCA for preliminary QC, thereafter upload imputation-server formatted files to publicly available imputation servers, and upon completion, users may follow through with both imputation and subsequent post-imputation analysis modules that are available within the RICOPILI framework[10]. Users are reminded that prior IRB approval, from their own institutions and/or institutions that provided access to the genotype data for the cohorts, might be necessary for using public genotype imputation services. For some cohorts with national laws preventing transfer of individual-level genotype data outside the country, it is not viable to impute to HRC at Michigan or Sanger as these services are US-based.

#### 4. Post-Imputation Module Description

RICOPILI conducts genome-wide association analysis using the imputed dosage files as input with PLINK (Purcell *et al*., 2007). Association testing of multiple cohorts can be performed in one step. Covariates can be added, and alternative phenotypes can be chosen. Meta-analysis can be performed either directly after genome-wide association across dosage files of multiple cohorts, or with RICOPILI formatted summary statistics. In the latter, summary statistics from multiple cohorts can be processed via the post-imputation module for meta-analysis. Meta-analysis is performed via METAL (Willer *et al.*, 2010). LD-clumped results ($R^2$ threshold of 0.1) are generated to identify genome-wide significant genomic regions, using a European reference as the default for clumping or another ancestry reference panel prepared for RICOPILI (note that reference panels could be built within the cluster, see section 5.1. Existing files for clumping purposes are also hosted here[11]). The left and right margin of the clumps is kept and partially or wholly overlapping clumping regions within less than 50kb distance are merged. Association analysis and meta-analysis are highly parallelized (*N* datasets x *N* genome chunk jobs). Outputs that are automatically generated in this module include:

- whole genome summary statistics with:
    - chromosome
    - rs-id
    - genomic position
    - allele 1 and 2
    - association values (odds ratio for allele 1, standard error, *P* value)
    - frequency information for cases and controls
    - quality score as observed variance divided by expected variance
    - number of cohorts for which the SNP was directly genotyped (vs imputed)
    - heterogeneity of effect test
    - raw and effective sample size per SNP
- annotated detailed Excel file of the LD-clumped regions with SNPs with $P < 0.0001$
- Manhattan plots in various forms (including for heterogeneity test)
- Q-Q plots
- region plots for genomic regions containing genome-wide significant SNPs
- forest plots depicting each genome-wide significant SNP effect by cohort
- QC reports for the meta-analysis and each cohort individually

[8] https://imputationserver.sph.umich.edu/index.html#!pages/home
[9] https://imputation.sanger.ac.uk/
[10] https://docs.google.com/document/d/18dupvU4kw11slREc1TUfwQwhO_eI0n_MeKVpwi4HLNA/ (Imputation server)
[11] https://sites.google.com/a/broadinstitute.org/ricopili/reference-panels

- standard LD score regression results
- additional summarizing files

As part of the RICOPILI post-imputation module, LD score regression (Bulik-Sullivan *et al.*, 2015) can be optionally performed to estimate the SNP-based heritability (SNP-*h*2) of the trait on the observed and liability scales, and to estimate the LD score intercept, which is informative about population stratification. Additional optional features of the post-imputation module include polygenic risk scoring (Purcell *et al.*, 2009) and conditional analysis which is informative for fine-mapping. The optional polygenic risk scoring command --bgscore allows prediction models to be tested in either i) independent target datasets available beyond those included in earlier summary statistics[12] or ii) the leave-one-out approach (See section 5.3) that allows each dataset included in the overall meta-analysis to serve as the target[13], while the remainder of the cohorts serve as the discovery sample, in each leave-one-out run.

A feature of the RICOPILI pipeline is that it permits the consolidation and generation of publication-ready figures and tables. As such, users may format meta-analysis summary statistics files that are compatible with RICOPILI and perform downstream visualization[14].

### 5. Additional Utility Modules

##### *5.1 Reference Builder*

If users are utilizing RICOPILI for the first time on their cluster, it will be necessary to build the reference files automatically used for pre-phasing, imputation, and annotations within the post-imputation module[15]. RICOPILI can build reference files from publicly available reference panels (e.g. 1000 genomes reference panel) or those available via permission (e.g. HRC).

##### *5.2 Replication*

Replication procedures are crucial to the robustness of GWAS results. RICOPILI also features the replicator module that allows replication of GWAS results where full summary statistics are unavailable for meta-analysis - single and multiple summary statistics are permitted for this replication module[16].

##### *5.3 Polygenic Risk Score Leave-One-Out Analysis*

If users have successfully completed imputation and post-imputation analysis and wish to perform polygenic risk score leave-one-out (LOO) analysis across all available cohorts, they will be able to carry this out within RICOPILI. This analysis leaves one cohort out at a time as the target for polygenic risk prediction analysis, while using all others as the training data.  The my.preploo2 module performs polygenic risk  score LOO analysis.

---

[12] https://docs.google.com/document/d/10jvLvnrPIz9zRKlRfMeW8HsHDxEIbs84yP3Ugjhj31E/ (PRS Scoring)
[13] https://docs.google.com/document/d/10UKUfX4wT9rP5zswH5SDR2Y1O2-wJVUw9mn4tSdaS-I/ (Leave-one-out scoring)
[14] https://docs.google.com/document/d/1o4bN_uLK4IEItXCSdeQkXfZwEpLuSCWlveJevRogi08/ (Postimp Module)
[15] https://docs.google.com/document/d/1pfJZmacumWHavtWP4aej2685c5gbLVA37rhba6M_vaw/ (Reference Builder)
[16] https://docs.google.com/document/d/1mvr2Kx2MrYaqlVIlE5nSpMePTO_qWkmSZySdT4AdBQw/ (Replication Module)

##### *5.4 Combining Data Across Distinct RICOPILI Projects*

In some cases, users might want to combine data across distinct RICOPILI imputations. This is especially so when one is interested in performing joint PCA across cohorts or polygenic risk score LOO analysis. The my.joinimp2 module allows and facilitates this process in a standardized manner.[17]

#### 6. HAPGEN Data Simulation Methods and Detailed Tutorial to Perform Quality Control, Imputation, Association Analysis and Meta-Analyses

To allow new users to familiarize themselves with RICOPILI and for experienced users to develop new functionality to the pipeline, we have simulated freely available GWAS data using HAPGEN (Su *et al.*, 2011). Given a reference panel of haplotypes, HAPGEN produces a sample of haplotypes with patterns of LD similar to those in the reference panel, in our case to 1000 Genomes Phase 3 data. Two datasets were prepared: one for a European ancestry population (EUR) (using 503 reference individuals), and another for an East Asian ancestry population (EAS) (using 504 reference individuals). After simulations, both datasets had 100,000 simulated "individuals" and around 80M variants. At the QC step, we removed variants with MAF < 0.005 and duplicated rs-ids and/or CHR:POS. After QC, 11,015,883 variants remained for the EUR dataset, and 9,952,334 variants remained for the EAS dataset. To create datasets that are appropriate for teaching and development purposes, we further modified the simulated datasets. From HAPGEN simulated data, we randomly selected 6000 samples from a European ancestry population (EUR) and 200 from an East Asian ancestry population (EAS) and extracted SNPs of a widely-used genotyping platform (Illumina OmniExpress). We then divided the EUR cohort into five subsets ($N$ = 2,000, 1,100, 1,000, 1,000, 1,000) and the EAScohort into two subsets (of $N$ = 100). We assigned case and control status randomly to each EUR cohort and assigned 2/98 and 5/95 cases/controls to EAS cohort 1 and 2, respectively. Finally, we merged the two EAS cohorts to two of the EUR cohorts. For teaching purposes, we introduced technical errors and population biases to these cohorts[18]. We selected cohort 1 and 2 to produce technical errors with false positive associations. After introducing population, technical and other association biases, we demonstrate the use of the RICOPILI pipeline to address them.

Our tutorial is available here in its most up-to-date form[19]. We also provide the tutorial in this Supplement after the reference list as an "Extended Data Analysis and User Guide". For new users of RICOPILI, it is important to note that each module in RICOPILI runs in its entirety using a single command line. This contributes to RICOPILIs ease-of-use and capability for assisting analysts to rapidly conduct GWAS, which becomes especially important in a consortia context when large amounts of data from many cohorts needs to be processed. The contents of our tutorial are as follows. We illustrate the pre-imputation/QC and PCA modules, and show how they can be used for rigorous QC and to help us visualize and better understand our data. While the pipeline automatically removes the problematic SNPs and samples from the analysis, we also illustrate the situations where we can manually remove problematic SNPs and samples if needed. After QC, we illustrate the imputation module to perform genotype imputation in the five cohorts using the HRC

---

[17] https://docs.google.com/document/d/1uD0VBFr-9UeALZ4rVssJcJG9855nQ7tgX6B-zoamosg/ (Postimp module)

[18] It is clear to us that these artificial biases do not reflect real world scenarios. But we calibrated them in such a way that the outcomes resemble classic values/plots of erroneous cohorts analysts have encountered in the past.

[19] https://docs.google.com/document/d/1ux_FbwnvSzaiBVEwgS7eWJoYlnc_o0YHFb07SPQsYjI/ (RICOPILI Tutorial)

reference panel. RICOPILI can also be used to prepare data to be deployed on Michigan and Sanger external servers for imputation. We illustrate the post-imputation module to perform GWAS and meta-analysis on imputed dosage data. We explore important results of meta-analysis, and illustrate how the post-imputation module can be used for downstream analysis after GWAS, such as polygenic risk scoring LOO analysis. An example of such a tutorial is available here[20].

#### 7. Cluster Portability

RICOPILI is portable to several LINUX-based high-performance computing (HPC) environments such as QSUB[21], BSUB[22], SLURM[23], and the Google Cloud Platform (GPC)[24]. Custom installation files for a number of specific HPC clusters including the Genetic Cluster Computer (LISA and Cartesius), Broad Institute HPC, LongLeaf HPC, and Northwell Health HPC are already available[25]. To allow users of other HPC clusters to setup and install RICOPILI we have provided comprehensive instructions[26]. Additional binaries and datafiles that could easily be unpacked from a tarball are provided, along with the pipeline, and can be downloaded using up-to-date download links within the installation document. For users who do not have access to HPC environments it would still be possible to use RICOPILI modules using the --serial --sepa combination. The additional switches allow RICOPILI to run on standalone machines with multiple CPUs. In cases where a HPC environment is not available, we recommend that users refer to the imputation module description (section 3), where computationally intensive pre-phasing and imputation procedures may be carried out on publicly available genomic imputation services and re-integrated into RICOPILI thereafter.

#### 8. RICOPILI Output Logging Stages

A useful feature for users of RICOPILI is the module log file system. While setting up the configuration file during installation of RICOPILI, users nominate a directory on their cluster that module log files are to be outputted to. Each module is complex, performing a variety of steps and tasks, yet is run in its entirety from a single command line the user submits. The command line calls the module that is being run, and specifies other parameters that depend on the module being run, such as input file names, output file names, QC operations and thresholds, reference populations, and covariate and phenotype input file names. Users can consult the log files that are automatically outputted from the the pipeline while it is running. This is especially helpful for tracking the progress of the pipeline and troubleshooting issues that contribute to unsuccessful module runs. Below we document the shorthand output that appears sequentially in the log file, so that users can understand the step that the pipeline is performing. The shorthand jobstep output in the log file is prepended by the path location from which the module was run and the submitted command line, and postpended is the date and time stamp that the pipeline started a jobstep. An example log entry for the PCA module is: "/home/user/depressionanalysis/usacohort/pca    pcaer --out usacohort --prefercase dep_usa_eur_user-qc.bim ref_EUR.bim ref_EAS.bim ref_SAS.bim ref_AMR.bim

---

[20] https://docs.google.com/document/d/1ux_FbwnvSzaiBVEwgS7eWJoYlnc_o0YHFb07SPQsYjI/ (RICOPILI Tutorial)
[21] https://docs.google.com/document/d/1oY5IA4a6yG_pmbvWJC8A6MTzjYoGzVlqQ_aXUwWCl8I/ (QSUB Installation)
[22] https://docs.google.com/document/d/1fNFnC3-rBZkmtH47Je_yUfGatB9qhDGi9HtMSA3_MPw/ (BSUB Installation)
[23] https://docs.google.com/document/d/1jNVAU0kGWZjSqUrshAkPIjs2YTnUwFG4-s0Ce18LE_g/ (SLURM installation)
[24] https://cloud.google.com/
[25] https://docs.google.com/spreadsheets/d/1LhNYIXhFi7yXBC17UkjI1KMzHhKYz0j2hwnJECBGZk4/ (Custom Install options)
[26] https://docs.google.com/document/d/14aa-oeT5hF541I8hHsDAL_42oyvlHRC5FWR7gir4xco/ (Installation module)

ref_AFR.bim  me300.000001     Thu_Apr__4_07:22:37_2019”. The shorthand for the pipeline step outputted in this log entry is “me300.00001”. Note that numbers after the period inform the user of the number of jobs sent to the cluster (i.e., jobstep.xx = xx jobs sent in parallel). We provide a succinct annotation below of each shorthand entry in the log in the sequential order that the user will see these appear for each module, including the hardware resources requested.

##### *8.1 Pre-Imputation/QC Module*

plague.1 : Platform guessing from .bim file; hardware resources: h_vmem=2g h_rt=1:00:00
qc.1 : Quality control/data management; hardware resources: h_vmem=2g h_rt=1:00:00
finished: confirmation that the module completed successfully

##### *8.2 Principal Components Analysis Module*

me300.000001 : Extracting 300 individuals for LD-pruning; hardware resources: h_vmem=3g -l h_rt=1:00:00
pruneprep.000001 : Preparation step for the LD-pruning; hardware resources: h_vmem=1g h_rt=1:00:00
prune1.000001 : First round of LD-pruning for the dataset; hardware resources: h_vmem=1g h_rt=1:00:00
prunebed.000001 : Prune plink file; hardware resources: h_vmem=1g h_rt=1:00:00
prune2.000001 : Second round of LD-pruning; hardware resources: h_vmem=1g h_rt=1:00:00
mepr.000001 : Merge all pruned datasets from the PCA; hardware resources: h_vmem=3g h_rt=2:00:00
genome.000002 : Creating the pi-hats –genome; hardware resources: h_vmem=4g h_rt=2:00:00
repov.000001 : Reporting for the pi-hat tables; hardware resources: h_vmem=1g h_rt=1:00:00
epca.000001 : Actual MDS/PCA creation with the eigenvectors; hardware resources: h_vmem=6g h_rt=2:00:00
asso.000020 : Genome-wide association of the PCs on the genotypes; 20 parallel jobs (20 PCs); hardware resources: h_vmem=2g h_rt=1:00:00
gwapl.000020 : Making manhattan plots from the association; hardware resources: h_vmem=2g h_rt=1:00:00
qqpl.000020 : Making qq-plots from the association; hardware resources: h_vmem=2g h_rt=1:00:00
lahu.000040 : Lambda hunting plots; hardware resources: h_vmem=2g h_rt=1:00:00
pcaplot.000001 : Rest of the PCA plots 1ds; 2ds etc; hardware resources: h_vmem=1g h_rt=1:00:00
finished:  confirmation that the module completed successfully

##### *8.3 Imputation Module*

buigue.000001: Build guessing, liftover to hg19 if necessary; hardware resources: h_vmem=2g h_rt_2:00:00
readref.000022 : Read reference for SNP information; hardware resources: h_vmem=2g h_rt=2:00:00

reresum.000001 : Sums up SNP information read from reference; hardware resources: h_vmem=1g h_rt=2:00:00
predep.000001 : preparation of datasets for deployment to imputation server; hardware resources h_vmem=2g h_rt=2:00:00
chepos.000001 : Alignment of position based on reference genomic position (Depending on 1KGP ref or HRC ref) – compares SNP names, reading rs names and converting from genotyping chips; hardware resources: h_vmem=8g h_rt=2:00:00
chefli.000001 : Checks and aligns SNP alleles based on the reference (1KGP/HRC) and resolves strandflips; hardware resources: h_vmem=8g h_rt=2:00:00
preph.000115 : Prephasing in genomic chunks; hardware resources: h_vmem=2g h_rt=2:00:00
pseudo.0005 : create pseudo case-controls for trios if trio dataset; hardware resources: h_vmem=13g h_rt=1:00:00
imp.000115 : Imputation; hardware resources: h_vmem=4g h_rt=2:00:00
dos.000115 : Additional step for reducing 3 dosages into 2 dosages (only for minimac); hardware resources: h_vmem=2g h_rt=2:00:00
dabg.000115 : Building best guess genotypes out of dosages based on genomic chunks; post-imputation QC procedures; hardware resources: h_vmem=2g h_rt=2:00:00
cobg_gw.000003 : Combining best guess genomic chunks into plink binary files, not filtered, moderately and strictly filtered; hardware resources: h_vmem=12g h_rt=2:00:00
clean.000008 : Removing intermediate/summarizing files; hardware resources: h_vmem=1g h_rt=2:00:00
du.000001 : Report on disk usage; hardware resources: h_vmem=1g h_rt=2:00:00
performer.00001 : Performance on each step; hardware resources: h_vmem=2g r_rt=2:00:00
vcf2dos_deploy.000115 : converting VCF files into dosage files; integrating imputation-server data for postimp module; hardware resources: h_vmem=4g h_rt=2:00:00
finished: confirmation that the module completed successfully

#### *8.4 Post-Imputation Module*

*8.4.1 Association Analysis*

daner.000114 : Pure association analysis on dosage chunks; hardware resources: h_vmem=1g; h_rt=2:00:00
dameta.000114 : Meta-analysis on the chunks; hardware resources: h_vmem=1g h_rt=1:00:00
damecat.000002 : Merging chunks into genome-wide files; hardware resources: h_vmem=1g h_rt=1:00:00
lth.000002 : Count SNPs and create files with different p-value thresholds are built e.g. p < 0.0001; p < 0.00001 etc; hardware resources: h_vmem=1g h_rt=1:00:00
areator.000002 : Clumping of top results; hardware resources: h_vmem=3g h_rt=1:00:00
areaplot.000075 : Plot region plots (independent loci); hardware resources: h_vmem=1g h_rt=1:00:00
forestplot.000077 : Plot forest plots (top index SNPs); hardware resources: h_vmem=1g h_rt=1:00:00
manhplot.000006 : Plot Manhattan plots; hardware resources: h_vmem=1g h_rt=1:00:00
qqplot.000002 : Plot QQ-plots; hardware resources: h_vmem=3g h_rt=1:00:00
lahu.000010 : Lambda hunting over different variations e.g. lambda/allele frequency; lambda/info score; hardware resources: h_vmem=2g h_rt=1:00:00
ldsc.000001 : Ldscore regression; hardware resources: h_vmem=4g h_rt=1:00:00

finished: confirmation that the module completed successfully

*8.4.2 Meta-Analysis*

chunk.000001 : Chunking genome-wide summary statistics into genomic chunks; hardware resources: h_vmem=1g h_rt=1:00:00
resmet.000293 : Meta-analysis of genome-wide chunks; hardware resources: h_vmem1g h_rt=1:00:00
damecat.000001 : Merging meta-analysis chunks into genome-wide files; hardware resources: h_vmem=1g h_rt=1:00:00
daner_het2p.000001 : If there were more than one files, these are the plots for the heterogeneity p-values; hardware resources: h_vmem=1g r_rt=1:00:00
lth.000004 : as above
areator.000004 : as above
areaplot.000127 : as above
areaplot.000098 : as above
forestplot.000233 : as above
manhplot.000012 : as above
qqplot.000004 : as above
ldsc.000001 : as above
lahu.000014 : as above
finished:  confirmation that the module completed successfully

*8.4.3 PRS Scoring*

danscore.000071 : scoring chunks
dsc.000001 : summarizing chunks and plotting
finished:  confirmation that the module completed successfully

*8.4.4 Clumping*

clump.000022 : plink clumping, but keeping information e.g. merging genes into the information
finished: confirmation that the module completed successfully

##### *8.5 Reference Builder Module (Optional)*

vcf_download.22 : download vcf file from reference panel with wget; 22 parallel jobs initiated; hardware resources: h_vmem=2g h_rt=2:00:00 (--walltimeplus)
bcftools_filtnorm.22 : filter on minor allele frequency counts; 22 parallel jobs initiated; hardware resources: h_vmem=2g h_rt=2:00:00 (--walltimeplus)
filter_filtnorm.22 : filters and reformats the vcf files from the above step; 22 parallel jobs initiated; hardware resources: h_vmem=2g h_rt=2:00:00 (--walltimeplus)
bcftools_impute2.22 : converts vcf into impute2 compatible format; 22 parallel jobs initiated; hardware resources: h_vmem=2g h_rt=2:00:00 (--walltimeplus)
vcf2plink.22 : convert to plink format from vcf; 22 parallel jobs initiated; hardware resources: h_vmem=2g h_rt=2:00:00 (--walltimeplus)
reformat.22 : rewrites legend files and bimfiles so that indels get coded as I and D and SNP names takes the form of “rs”; 22 parallel jobs initiated; hardware resources: h_vmem=2g h_rt=2:00:00 (--walltimeplus)
trans_filtnorm.22 : Rewrites filtnorm with legend changes from refdir_navi2; 22 parallel jobs initiated; hardware resources: h_vmem=2g h_rt=2:00:00 (--walltimeplus)
prep_minimac_bgz.22 : Creates FILE.vcf.bgz (a block-gzipped VCF file) for eagle and minimac3; 22 parallel jobs initiated; hardware resources: h_vmem=2g h_rt=2:00:00 (--walltimeplus)
prep_minimac_tabix.22 : Creates FILE.vcf.bgz.tbi (a tabix index) for eagle and minimac3; 22 parallel jobs initiated; hardware resources: h_vmem=2g h_rt=2:00:00 (--walltimeplus)
prep_minimac_m3vcf.22 : Creates FILE.m3vcf.gz (a Minimac3 version of the vcf file); 22 parallel jobs initiated; hardware resources: h_vmem=24g h_rt=02:00:00 (--walltimeplus)
vcf2bcf.22 : converts vcf to bcf, improves efficiency of prephasing by a lot also tabix is used to index the resulting file afterwards; hardware resources: h_vmem=2g h_rt=2:00:00 (--walltimeplus)
chunk.132 : Creating a meta-file that contain chunk sizes for legend files; 132 parallel jobs; hardware resources h_vmem=2g h_rt=2:00:00
pobed.132 : Creates population specific plink binaries for each chromosome (for LD pruning jobs); hardware resources: h_vmem=4g h_rt=2:00:00 (--walltimeplus)
finished: confirmation that the module completed successfully

**Note: for full explanation of RICOPILI file output structure and other helpful tutorials please visit** [https://sites.google.com/a/broadinstitute.org/ricopili/home](https://sites.google.com/a/broadinstitute.org/ricopili/home)

**Supplementary Table 1: Available GWAS pipelines**

| Name | Free for Implementation | Scalability | User Flexibility (User controls analysis parameters) | Extensiveness of Pipeline | Platform/Language | Software/Libraries |
|---|---|---|---|---|---|---|
| GWAS-Pipeline | No | Cluster based (Private)<br>*Note: Not easily implemented on other clusters - not open scourced* | Yes | QC: In-house scripts<br>Imputation: Unknown<br>PCA: In-house scripts<br>GWAS Association: Plink<br>Polygenic Prediction: Not carried out<br>Visualization: Manhattan, region, qqplots | Linux; Python; R | Plink |
| GWASPi | Yes | Desktop | Yes | QC: Software Specific<br>Imputation : Not carried out<br>PCA: Not carried out<br>GWAS Association : Software Specific<br>Visualization: Manhattan, qqplots | Windows; Java | JavaDB SQL, NetCDF-3 etc |
| Imputation Servers (e.g. Michigan, Wellcome Trust | Yes | Cluster based (Cloud)<br>*Requires specific IRB permissions for use* | No | QC: Not part of pipeline<br>Imputation: Multiple Reference Panels (e.g. HRC, 1000 genomes, HAPMAP, CAAPA), Multiple pre-phase tools, Imputes on Minimac3<br>GWAS: Association: Not Carried out<br>Visualization: Not Carried out | Linux | Eagle2, Shapeit, Hap-UR Minimac3 |
| Gimpute | Yes | Desktop/Cluster<br>*Note: Serial input with single multi-core machine* | Yes | QC: qctool or gtool<br>Imputation : Shapeit/Impute<br>PCA: GCTA<br>GWAS Association : Not carried out<br>Visualization: Not carried out | Linux; R | Shapeit, Impute4, Plink, GCTA |
| RICOPILI: Rapid Consortias Imputation Pipeline | Yes | Cluster (Private) - Open sourced<br>Desktop (Using multi-core processing)<br>*Flexible cross platform implementation QSUB,BSUB,SLURM* | Yes | QC: Integrated with Plink (Standardized reporting protocol) - also handles Trio dataset<br>Imputation: *Flexible* (Pre-phase, Shapeit, or Eagle2), Imputation (Impute4 or Minimac3), Reference Panels (1000 genomes or HRC)<br>PCA : Eigenstrat<br>GWAS Association: Plink<br>Meta-Analysis: Metal<br>Polygenic prediction: Software Specific Scripts<br>Visualizations: Manhattan, Region, QQ plots, as well as PRS prediction plots. Other diagnostic visualizations are also provided | Linux; Perl; R | Eagle2, Shapeit, Impute4 Minimac3, Plink, LDSC, Eigenstrat |

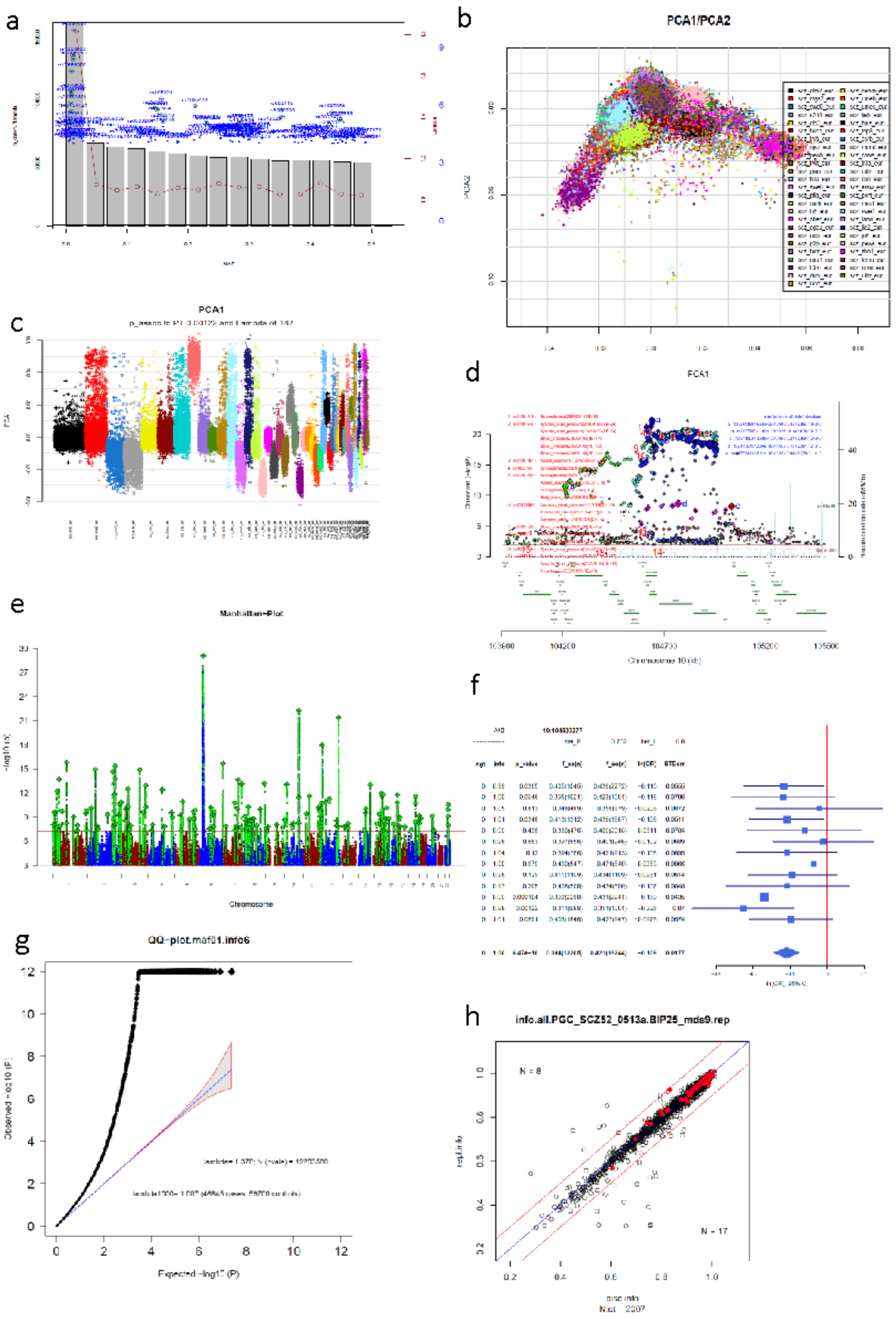


**Supplementary Figure 1. RICOPILI visualizations**

*Note:* a. Quality control filtering; b. 2-dimension PCA visualization; c. 1-dimension PCA visualization; d. Region plot; e. Manhattan plot; f. Forest plot; g. QQ-plot; h. replication plot

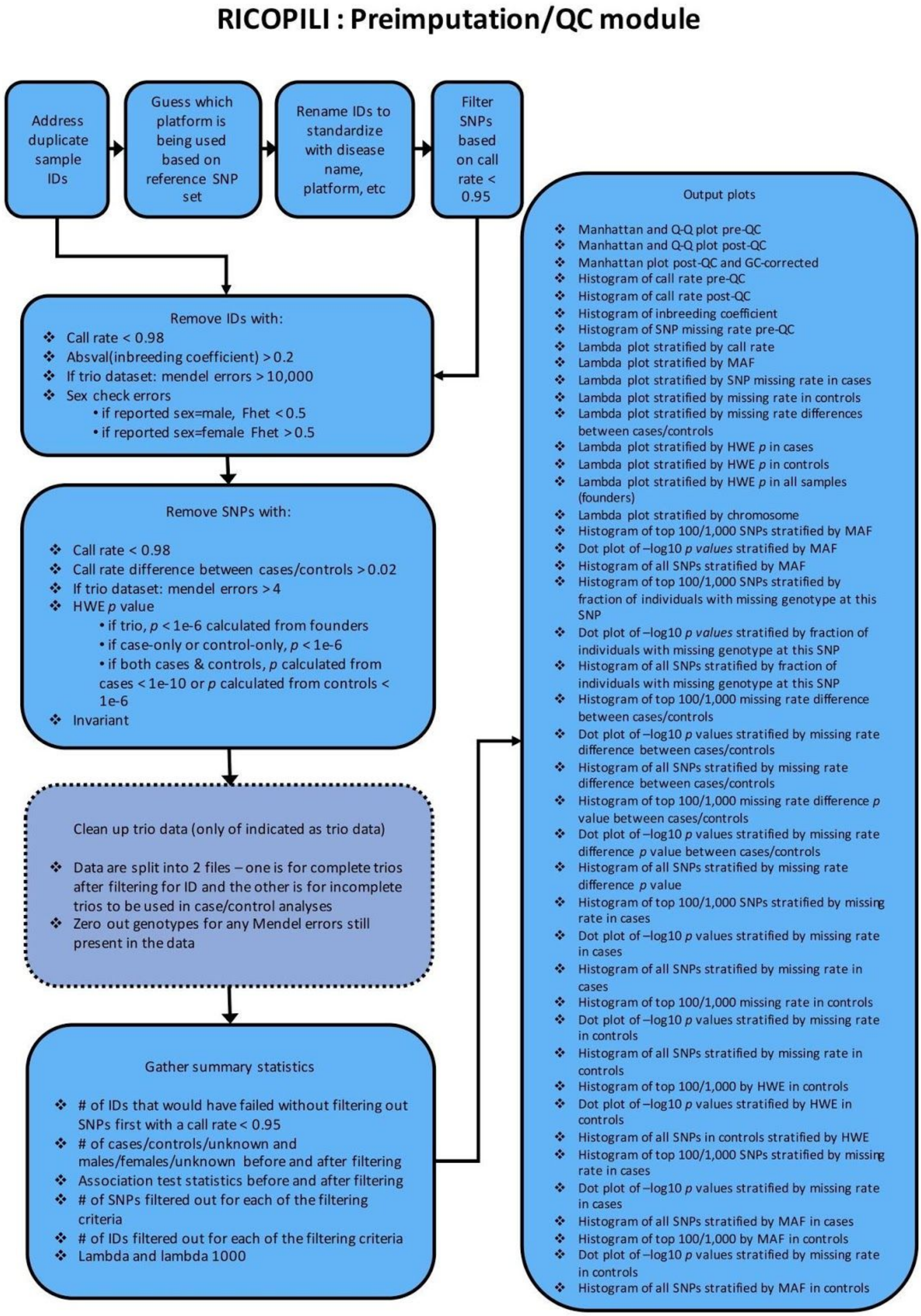


**Supplementary Figure 2. Pre-imputation/quality control module workflow**

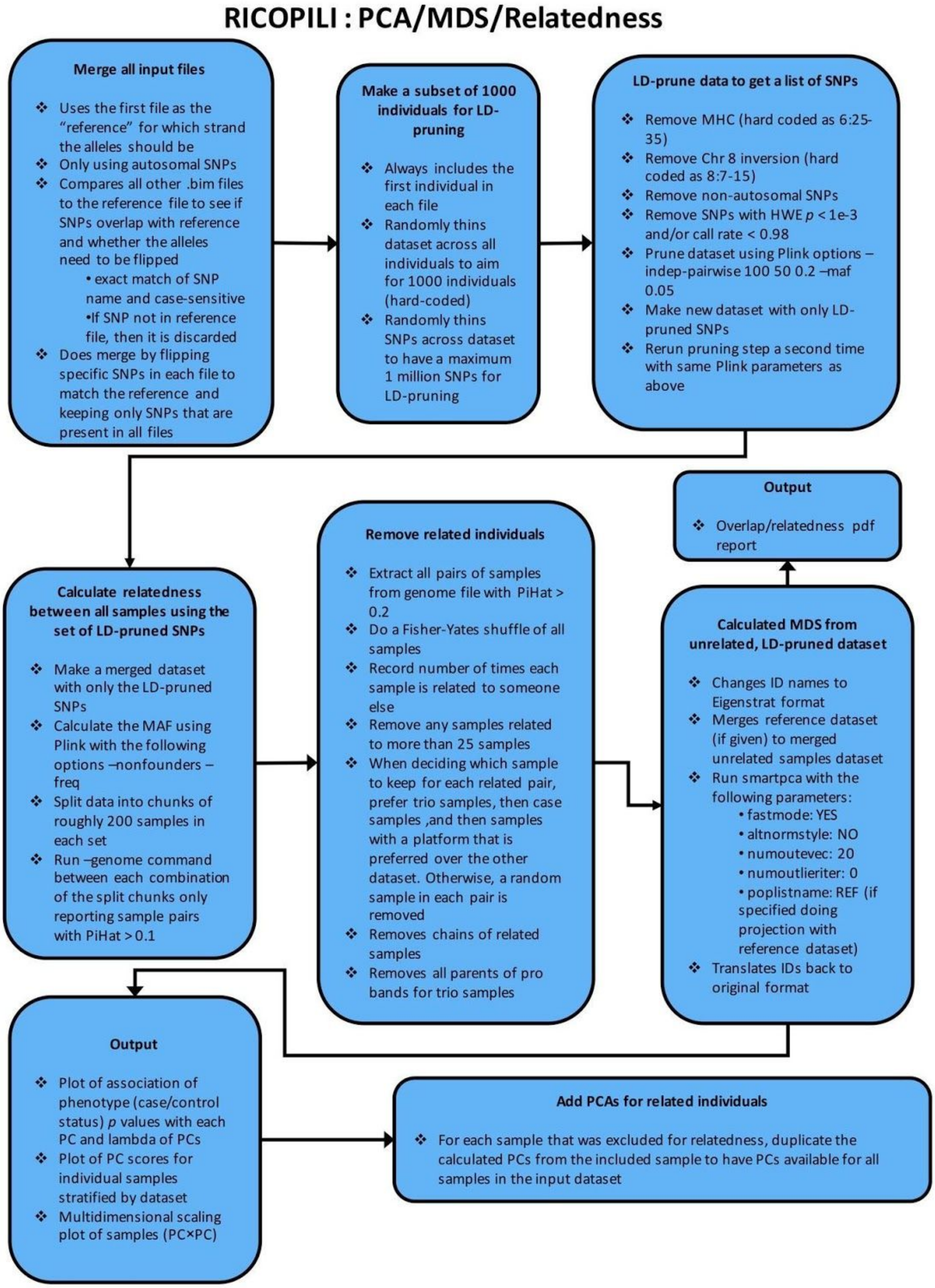


**Supplementary Figure 3. Principal Components Analysis (PCA) module workflow**

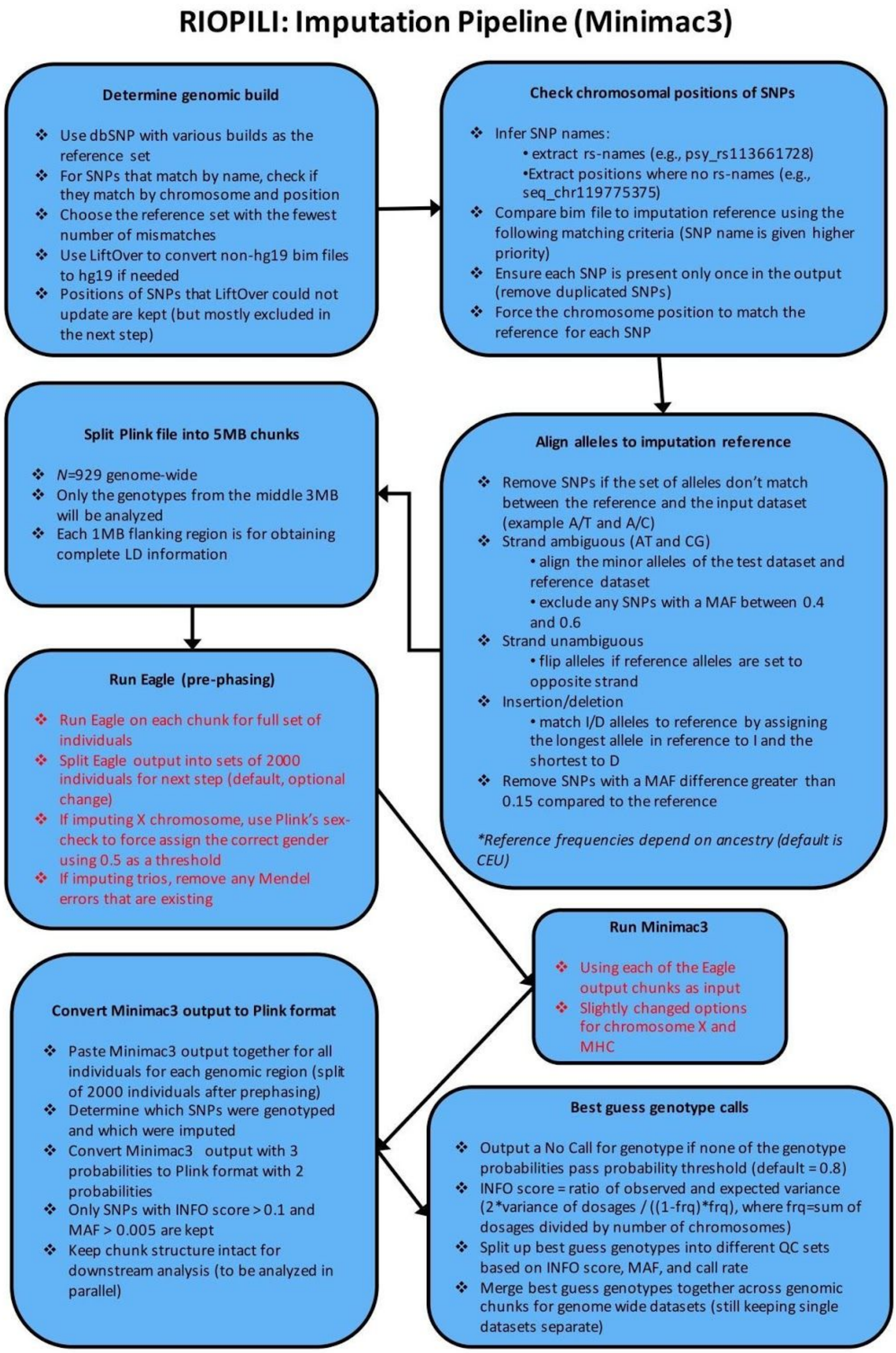


**Supplementary Figure 4. Imputation module workflow**

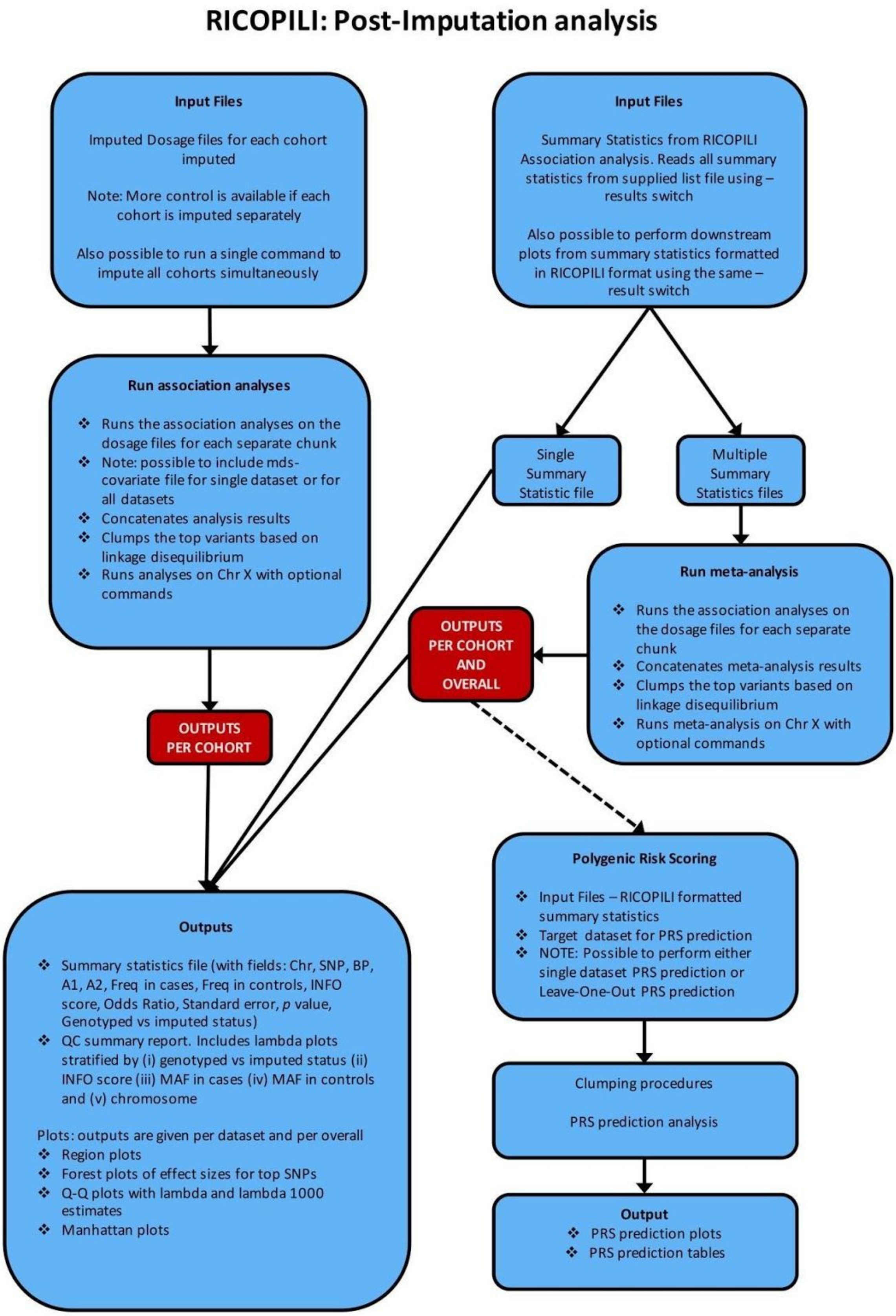


**Supplementary Figure 5. Post-Imputation module workflow**

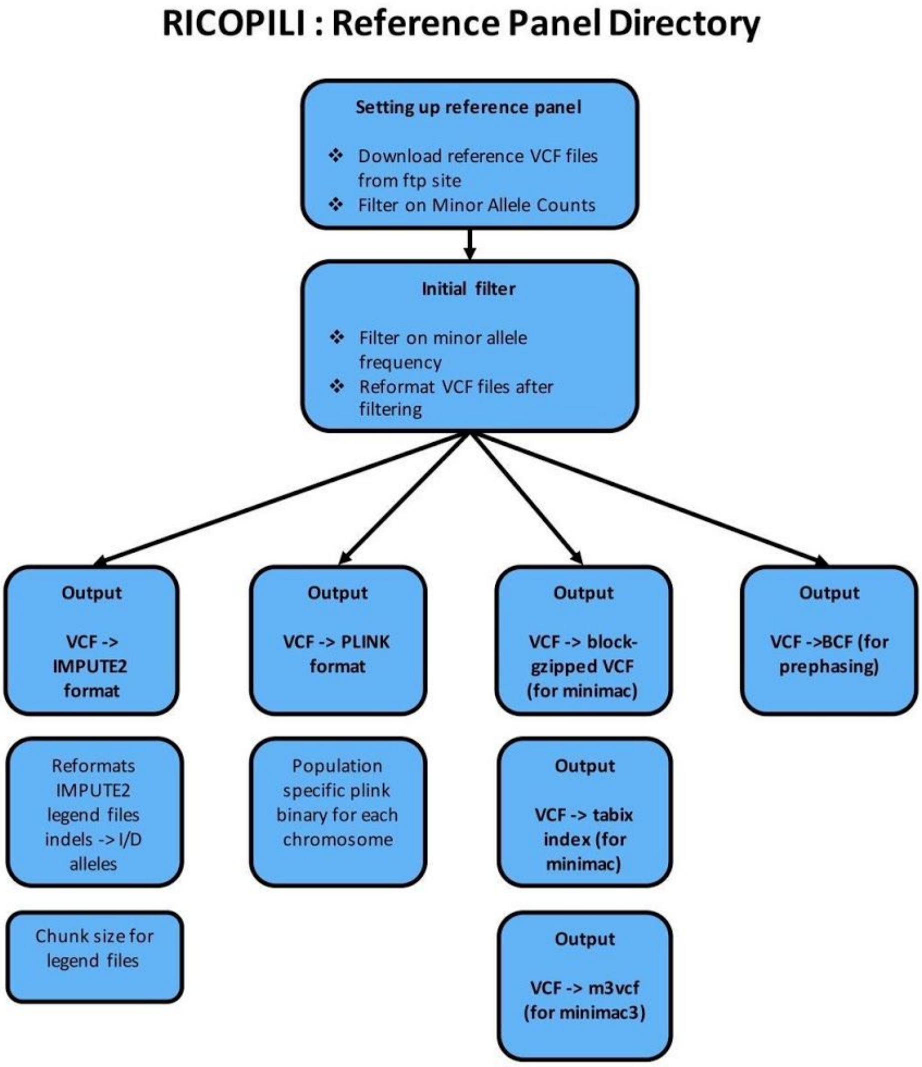


**Supplementary Figure 6. Reference builder module workflow**

**Websites**

**RICOPILI FAQ/Download/Tutorial** https://sites.google.com/a/broadinstitute.org/ricopili/home
**RICOPILI Github** https://github.com/Nealelab/ricopili
**LiftOver** https://genome.sph.umich.edu/wiki/LiftOver
**Eigenstrat** https://www.hsph.harvard.edu/alkes-price/software/
**PLINK** http://www.cog-genomics.org/plink2
**EAGLE** https://www.hsph.harvard.edu/alkes-price/software/
**Minimac3** https://genome.sph.umich.edu/wiki/Minimac3
**METAL** https://genome.sph.umich.edu/wiki/METAL_Documentation

### Tutorial: Extended Data Analysis and User Guide

To further help users understand the basic data analytic procedures of the RICOPILI pipeline, we have included a tutorial, the Extended Data Analysis and User Guide, that covers a step-by-step showcase of the simulated data, described in Section 6 of the Supplementary Information, and output of data analysis conducted via RICOPILI.

##### *Cohort descriptions*

From Hapgen simulated data we randomly selected 6,000 european (EUR) and 200 asian (ASN) individuals and extracted the SNPs of a widely-used genotyping platform (Illumina OmniExpress). We then divided the EUR cohort into five subsets (*N* = 2,000, 1,000, 1,000, 1,000, 1,000) and the ASN cohort into two subsets (*N* = 100 each). We assigned case and control status randomly to each EUR cohort and assigned 2/98 and 5/95 cases/controls to the two ASN cohorts. Finally, we merged the two ASN cohorts to EUR cohorts 1 and 3. The composition of the resulting five cohorts is described in Extended Data Table 1. The cohorts are in PLINK binary format.

| Cohort # | datasets | n | cases | controls | SNPs | interventions |
|---|---|---|---|---|---|---|
| 1 | sim_sim1a_eur_sa_merge.miss | 2,100 | 985 | 1,115 | 547,764[27] | Introduced population stratification (by merging it with asian cohort 1) and also technical errors |
| 2 | sim_sim2a_eur_sa_merge.miss | 1,000 | 474 | 526 | 593,970 | Only technical errors |
| 3 | hapgen_sample3a | 1,100 | 478 | 622 | 547,764[1] | Only population stratification by merging with asian cohort . |
| 4 | hapgen_sample4b | 1,000 | 483 | 517 | 593,970 | Shares 10 overlapping individuals with cohort 5 |
| 5 | hapgen_sample5a.ph | 1,000 | 516 | 484 | 593,970 | - |

Extended Data Table 1: Cohort descriptions

##### *Technical interventions and biases introduction*

To demonstrate the functionality of RICOPILI, we introduced technical errors and association biases to these cohorts. We selected cohort 1 and 2 to produce technical errors with false positive associations. In order to do so, we performed the following steps **separately for cases and controls** in both of the cohorts[28].

[27] The number of SNPs are less than the other cohorts as these were formed by a merging asian and european cohorts, so we only took the overlapping SNPs between them.
[28] It is clear to us that these artificial biases do not reflect real world scenarios. But we calibrated them in a way that outcome resembles classic values/plots of erroneous cohorts we encountered in the past.

a. To create autosomal heterozygosity rate deviations in individuals, we changed the heterozygous genotypes to homozygous for all SNPs in 10 selected probands.
b. To create missingness per individual, a set of 100 probands was selected, and a missing rate of SNPs between 0-10% (from a skewed distribution with higher probability of low missing rates) was introduced.
c. To create sex errors, we randomly selected 10 (male/female) and swapped their gender assignments.
d. To create missing SNPs, we randomly selected 2% SNPs of all individuals and introduced missing genotypes by choosing missingness rates between 0-10% (form a skewed distribution with higher probability of low missing rates).
e. To create Hardy Weinberg disequilibrium per SNPs, we randomly selected 2% of all SNPs and introduced an artificial excess of homozygosity.
f. To create false positive associations SNPs, we selected 20 SNPs and flipped allele1 and allele2 while introducing missingness in these SNPs.
g. Finally, cases and controls were merged back into a single cohort.

##### *Installation*

First, install the latest version of the RICOPILI pipeline on your High Performance Computer(HPC). This document here[29] describes the process of custom installation in detail.

##### *Download data*

The genotype data described in Extended Data Table 1 can be downloaded from here: https://personal.broadinstitute.org/sawasthi/share_links/afABqiK7luSzPwFjag1g5ij8ySlabj_gwas-qcerrors.py/

##### *Quality control*

These two modules of the pipeline are used for quality control.

- Pre-imputation/QC module[30] : for technical quality control of SNPs and individuals.
- PCA module[31] : for genomic quality control (ancestry checks, overlapping/related individuals).

Here, we introduce the functions of the Pre-imputation/QC module and PCA module to prepare genotyped datasets for genome-wide data analysis. The Pre-imputation/QC module performs standard GWAS quality control procedures for stipulated datasets. We will demonstrate in the ensuing sections multiple quality control runs, iteratively cleaning up the five HAPGEN datasets, until they are appropriate for downstream imputation and association analysis.

First, we placed all the above cohorts (Extended Data Table 1) into an empty directory and started the Pre-imputation/QC module using the command below from within that directory.

```
-bash:~$> preimp_dir --dis sim --out hapgen_5cohorts
```

[29] https://docs.google.com/document/d/14aa-oeT5hF541I8hHsDAL_42oyvlHRC5FWR7gir4xco/
[30] https://sites.google.com/a/broadinstitute.org/ricopili/preimputation-qc
[31] https://sites.google.com/a/broadinstitute.org/ricopili/pca

The *'preimp'* command calls this module. The command line  shown will run the module with default QC parameters, which are described in the earlier sections (Supplementary Information, Section 1) Flag *'--dis'* takes an abbreviation for the study phenotype, please use three characters[32] (we used "sim" for "simulated data"). After the first invocation of the command, users will be prompted to fill out *'[--dis].names'* (in this case it will be *'sim.names'*; Extended Data Figure 1).[33] Users are advised to use informative study names (please read recommendations in the comments) for each of the five cohorts under the column 'STUDYNAME' and leave "0" under column 'EXCLUDE' to keep these datasets in this QC run. Mark this first QC with "1" in the 'QCCYCLE' column.

```
###STUDYNAME: 5 alphanumeric characters, recommeded city or study abbreviation and wave, e.g. lond1, cloz2
###              please negotiate with PIs about this name
###BFILE:      bed-filename root, e.g. if you dataset is called YOURDATA.bed, then please use YOURDATA in this column
###QCCYCLE:    numeric indicating the rounds of quality controls you have already performed
###              this will be appended to the resulting files with -qc1 or -qc2
###EXCLUDE:    set 0 if you want this dataset being included in the preimp process
STUDYNAME     BFILE                     QCCYCLE        EXCLUDE
hap5a         hapgen_sample5a.ph            1              0
hap4a         hapgen_sample4b               1              0
hap3a         hapgen_sample3a               1              0
hap1a         sim_sim1a_eur_sa_merge.miss   1              0
hap2a         sim_sim2a_eur_sa_merge.miss   1              0
```

Extended Data Figure 1: Screenshot '*sim.names*' (*[--dis].names*) from first '*preimp*'.

Save *'[--dis].names'* and repeat the command below in the same directory.

```
-bash:~$> preimp_dir --dis sim --out hapgen_5cohorts
```

A successful job submission will end with the message in Extended Data Figure 2.

```
------------------------------------------------------------
1 jobs successfully submitted
please see tail of /home/unix/sawasthi//preimp_dir_info for regular updates
also check bjobs -w for running jobs
possibly diffent command on different computer cluster: e.g. qstat -u USER
you will be informed via email if errors or successes occur
------------------------------------------------------------
```

Extended Data Figure 2: Message of successful *'preimp'* jobs submissions

RICOPILI users are informed via email after successful completion of the module, if the email program is properly installed on your system. Nevertheless, it is recommended to check running modules on your HPC in the file *preimp_dir_info*[34] (Extended Figure 3; which is in your home directory or in the *loloc*-directory you specified in the configuration file during custom installation). After successful completion of the module, a detailed QC report for each cohort is found in the subdirectory *'qc/'* (Extended Data Table 2). We will now describe the results from the module and

---

[32] Three character names for some disorder/phenotypes used are here.

[33] We assume you are familiar with the terminal editors "vim" or "emacs". If not, please refer to these tutorials (https://vim.fandom.com/wiki/Tutorial , http://www.gnu.org/software/emacs/manual/html_node/emacs/index.html).

[34] Similar (*'preimp_dir_info', 'pcaer_info', 'pcaer_info', 'impute_dir_info', 'postimp_navi_info', 'replicator_info'*) log files exist for each RICOPILI module in the user's home directory or in the *loloc*-directory specified during custom installation. These logfiles record the job step of running processes and can be used to search for  locations and commands from earlier runs, and to troubleshoot unsuccessful module runs. It is recommended not to delete these files and to make regular backup copies.

will also perform further QC action if necessary on each cohort separately. It is recommended that users start a new directory using the *mkdir* command for each QC run.

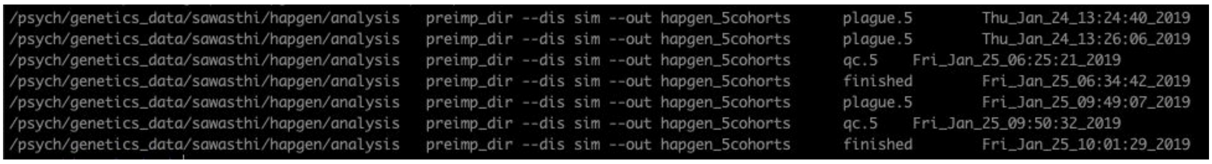


Extended Data Figure 3: Screenshot of *'preimp_info'*

| **Cohort #** | **Pre-QC'd genotype file names** | **QC report files names** | **Post-QC'd genotype file names** |
|---|---|---|---|
| 1 | sim_sim1a_eur_sa_merge.miss | sim_hap1a_eur_sa-qc1.pdf | sim_hap1a_eur_sa-qc1 |
| 2 | sim_sim2a_eur_sa_merge.miss | sim_hap2a_eur_sa-qc1.pdf | sim_hap2a_eur_sa-qc1 |
| 3 | hapgen_sample3a | sim_hap3a_eur_sa-qc1.pdf | sim_hap3a_eur_sa-qc1 |
| 4 | hapgen_sample4b | sim_hap4a_eur_sa-qc1.pdf | sim_hap4a_eur_sa-qc1 |
| 5 | hapgen_sample5a.ph | sim_hap5a_eur_sa-qc1.pdf | sim_hap5a_eur_sa-qc1 |

Extended Data Table 2: List of file names of (pdf) QC reports and pre/post-QC'd file names of each cohort from the first Pre-imputation/QC module run.

Each file name (QC'd genotypes and report pdfs in Extended Data Table 2) starts with the three character (abbreviation of the study phenotype; "sim" in this case) argument given to the *'--dis'* flag in the *'preimp'* command line. This is followed by the study name as filled in *'[--dis].names'* under the column 'STUDYNAME', then the population, which is European, designated as "eur" (population is set to european by default. Users may change this by using *--popname* in *'preimp'*). Finally the initials of the analyst are included in the output filename (here "sa"), which are indicated during the RICOPILI installation procedure [35]. The 'qc1' suffix in the file names implies that these belong to the first cycle of the *'preimp'* module run.

[35] Your file names will be almost the same as presented in Extended Data Table 2, or later tables in this tutorial, except that the "sa" (initials of the analyst) will be replaced with yours or the initials that were fixed during RICOPILI /installation.

#### Cohort 1 (sim_sim1a_eur_sa_merge.mis.bed/.bim/.fam)

This cohort consists of 983 cases, 1,117 controls of european population and 2 cases and 98 controls from asian population including technical biases for SNPs as described in sections 1 and 2.

#### Preimp/QC results:

**File: sim_hap1a_eur_sa-qc1.pdf**

### 1 General Info

#### 1.1 Size of Sample

| Test | pre QC | post QC | exclusion-N |
|---|---|---|---|
| Cases,Controls,Missing | 985,1115,0 | 929,1059,0 | 56,56,0 |
| Males,Females,Unspec | 1332,768,0 | 1262,726,0 | 70,42,0 |
| SNPs | 547764 | 507493 | 40271 (7.4%) |

#### 1.2 Exclusion overview

- would have excluded 73 individuals without pre-filter (SNP-Missing 0.05)

| Filter | N |
|---|---|
| SNPs: call rate < 0.950 (pre - filter) | 107 (0.0%) |
| IDs: call rate (cases/controls) < 0.980 | 73 (36/37) |
| IDs: FHET outside +- 0.20 (cases/controls) | 20 (10/10) |
| IDs: Sex violations -excluded- (N-tested) | 26 (2100) |
| IDs: Sex warnings (undefined phenotype / ambiguous genotypes) | 1 (0/1) |
| SNPs: call rate < 0.980 | 8051 (1.5%) |
| SNPs: missing difference > 0.020 | 5472 (1.0%) |
| SNPs: without valid association p-value (invariant) | 0 (0.0%) |
| SNPs: HWE-controls < 1e-06 | 16427 (3.0%) |
| SNPs: HWE-cases < 1e-10 | 10818 (2.0%) |
| Warning: genomewide significant SNPs (autosomal/known) | 16 (16/0) |

Extended Data Figure 4: Detailed QC report of first *'preimp'* on cohort 1

This QC report (Extended Data Figure 4) show most of the artificial technical errors flagged and rectified by RICOPILI. 112 individuals were excluded due to low call rate, sex violation and out of bounds heterozygosity deviation rates[36]. 40,271 (7.4%) SNPs were excluded due to high HWE deviation, and high missing rates. After QC we were left with 507,493 which would be adequate for proper imputation procedures (rough minimum ~250K). However, we noted that there were 16 SNPs significantly associated with the phenotype even after GC correction (Genomic Control for lambda inflation). This appeared to be a red flag. False positives would carry though imputation and should be accounted for before prior to implementing computationally expensive procedures. QQ plots, both pre and post QC (Extended Data Figures 5 and 6), show highly inflated lambda values, which did not appear to be resolved during technical quality . These are likely driven by population stratification (two genetically distinct population (eur/asn) in this cohort) or sample-overlap.

[36] In real world scenario 112 could be too many individuals to lose. Thus it make sense to double check your genotype data especially given this high number of sex errors. We do not inspect the data as these errors are artificially introduced.

#### 2 Manhattan

##### 2.1 Manhattan-Plot - pre-QC (QQplot with MAF 0.02)

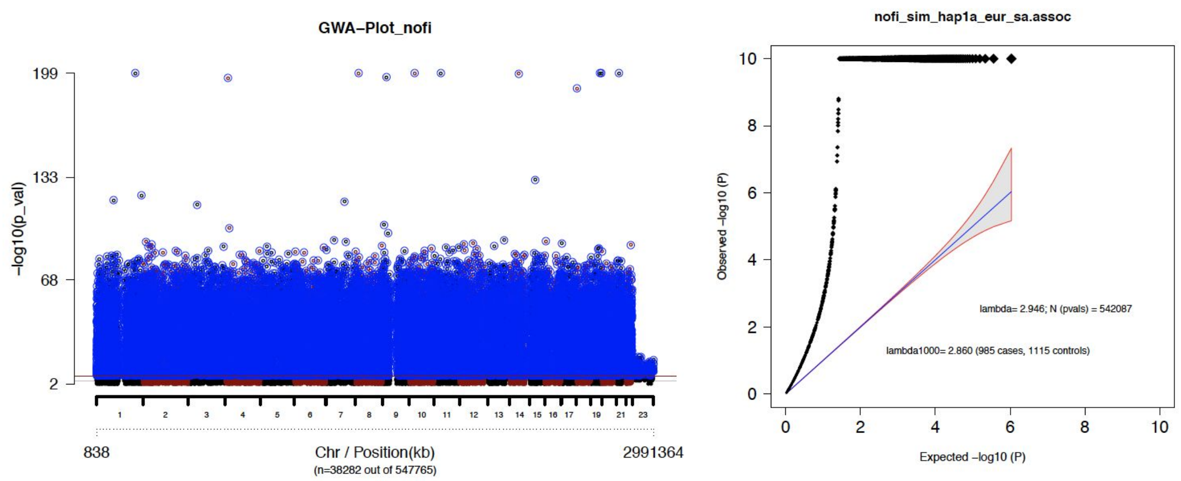


Extended Data Figure 5: Pre-QC QQ plot from first *'preimp'* on cohort 1

##### 2.2 Manhattan-Plot - post-QC (QQplot with MAF 0.02)

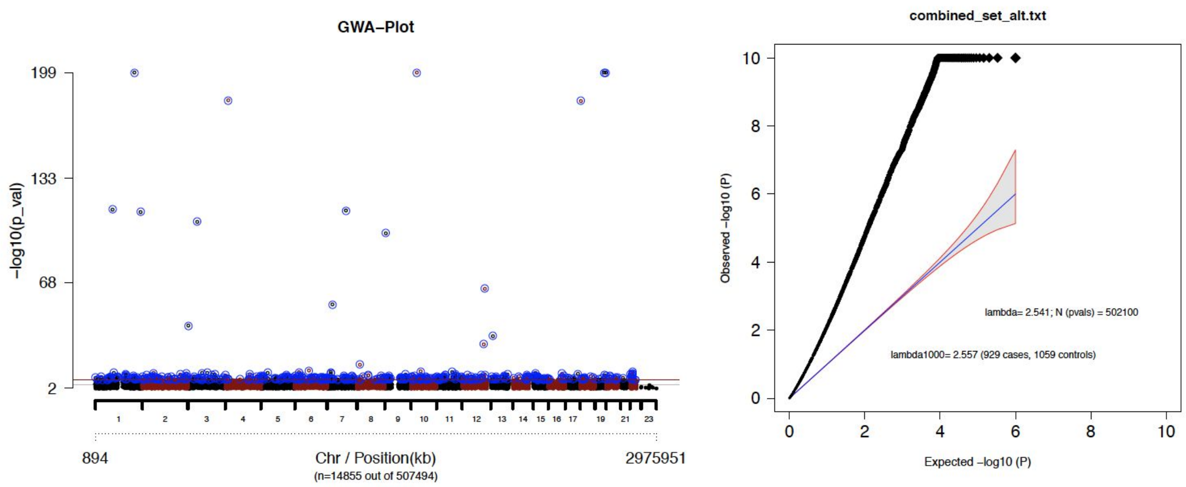


Extended Data Figure 6: Post-QC QQ plot from first *'preimp'* on cohort 1

We used the '*pcaer*' module to address population stratification and sample overlap. With this module principal component analysis (PCA) was carried out on the post-QC'd data **(sim_hap1a_eur_sa-qc1.bed/.bim/fam; Table 2)** to identify and visualize population structures in the data as well as visualizing sample-overlap and relatedness.

#### Principal component analysis

To start the *'pcaer' module we created symbolic links of QC-ed files into an empty directory and* ran the following command:

```
-bash:~> pcaer --prefercase --preferfam --out pca_hap1a
sim_hap1a_eur_sa-qc1.bim
```

Users would get a successful job submission message similar to that in the earlier *'preimp'* module. On completion of the analysis, an email from the system would be sent to the user. It is always advisable to check regularly *'pcaer_info'* in the home directory (or the home directory specified in the *'loloc'* file). A "finished" entry will indicate a completed analysis.

**File: pca_hap1a.menv.mds.2d.pdf.gz**[37]

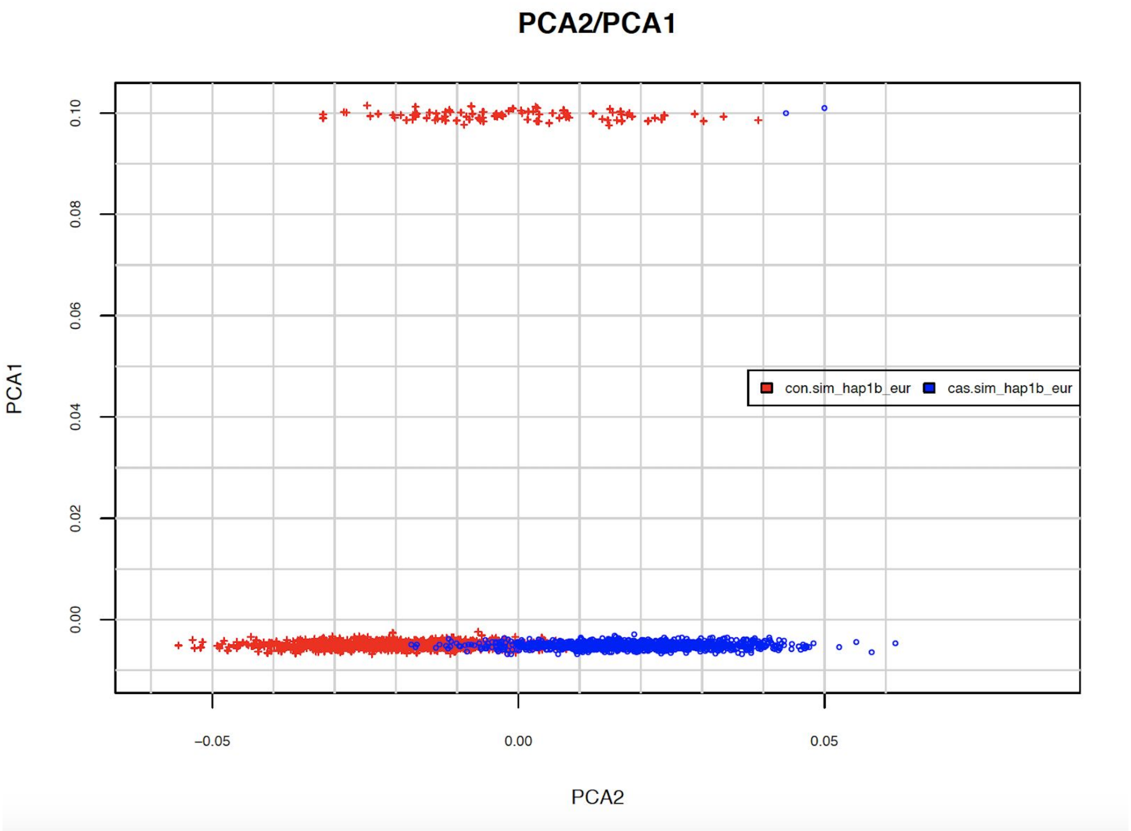


Extended Data Figure 7: PCA plot for cohort 1

Extended Data Figure 7 shows the visualization of the first two principal components where two distinct populations were identified. This is consistent with the merging of the European and East Asian ancestry samples where we introduced population stratification and technical errors to the genotyped datasets at the beginning of this section. To restrict analysis to only the European ancestry, we chose a threshold that will exclude the East Asian subset (e.g. PCA1=0.01).[38] Users may use the **pca_hap1a.menv.mds**[39] file for the purpose of filtering population outliers. The file also lists the value of all 20 PCs (principal components) of each individual.

---

[37] All the files mentioned in the '*pcaer*' module are found in its working directory.
[38] Of course these threshold depend on the data, some researchers also want to apply more sophisticated algorithms with euclidean distance.
[39] Since this file contains a non-overlapping subset of individuals we want to use it for including individuals (rather than excluding individuals on the "other side" of the threshold).

```
-bash:~> awk '$4<0.01' pca_hap1a.menv.mds >
hap1a.eur.sample.txt
```

Aside from identifying population stratification or outliers the *'pcaer'* module checks for overlapping/related individuals. To identify if there were overlapping individuals (within the cohort) we checked the **pca_hap1a.mepr.overlap** ([--out].mepr.overlap) file which is included in the following tarball **pca_hap1a.menv.mds.tar.gz.**

```
-bash:~> tar -xvzf pca_hapa.menv.mds.tar.gz
```

The file **[--out].mepr.overlap** lists all the overlapping/related individuals with PIHAT values greater than 0.2. Here **pca_hap1a.mepr.overlap** is empty indicating there are no overlapping/related individuals. Detailed analysis that identifies overlapping individuals will be discussed in later sections.

Individuals with European ancestry from the file **'hap1a.eur.sample.txt'**, were extracted using plink --keep. Such that a new set of genotype files could be created

```
-bash:~> plink --bfile sim_hap1a_eur_sa-qc1 --keep
hap1a.eur.sample.txt --make-bed --out hap1a_only_eur
```

These files were copied in a new empty directory, where we performed *'preimp'* on the second version of the data i.e. without the Asian individuals **(hap1a_only_eur.bed/.bim/.fam)** using similar command as described above.

```
-bash:~> preimp_dir --dis sim --out hap1a_eur
```

We altered '*sim.names*' (*[--dis].names*) to "hap1b" under STUDYNAME and "2" under QCCYCLE (Extended Data Figure 8) since this was the second cycle of '*preimp*' on cohort 1.

```
###STUDYNAME: 5 alphanumeric characters, recommeded city or study abbreviation and wave, e.g. lond1, cloz2
###              please negotiate with PIs about this name
###BFILE:     bed-filename root, e.g. if you dataset is called YOURDATA.bed, then please use YOURDATA in this column
###QCCYCLE:   numeric indicating the rounds of quality controls you have already performed
###              this will be appended to the resulting files with -qc1 or -qc2
###EXCLUDE:   set 0 if you want this dataset being included in the preimp process
STUDYNAME      BFILE           QCCYCLE    EXCLUDE
hap1b          hap1a_only_eur     2          0
```

Extended Data Figure 8: Screenshot '*sim.names*' (*[--dis].names*) from second '*preimp*' on cohort 1.

We saved *'[--dis].names'* and repeated the command below in the same directory.

```
-bash:~> preimp_dir --dis sim --out hap1a_eur
```

| S.No | Pre-Qc’d genotype file names | QC reports files names | Post-Qc’d genotype file names |
|---|---|---|---|
| 1 | hap1a_only_eur | sim_hap1b_eur_sa-qc2.pdf | sim_hap1b_eur_sa-qc2 |

Extended Data Table 3: Pre/Post QC’d files from second *‘preimp’* on cohort 1.

##### File: sim_hap1b_eur_sa-qc2.pdf

2.2 Manhattan-Plot - post-QC (QQplot with MAF 0.02)

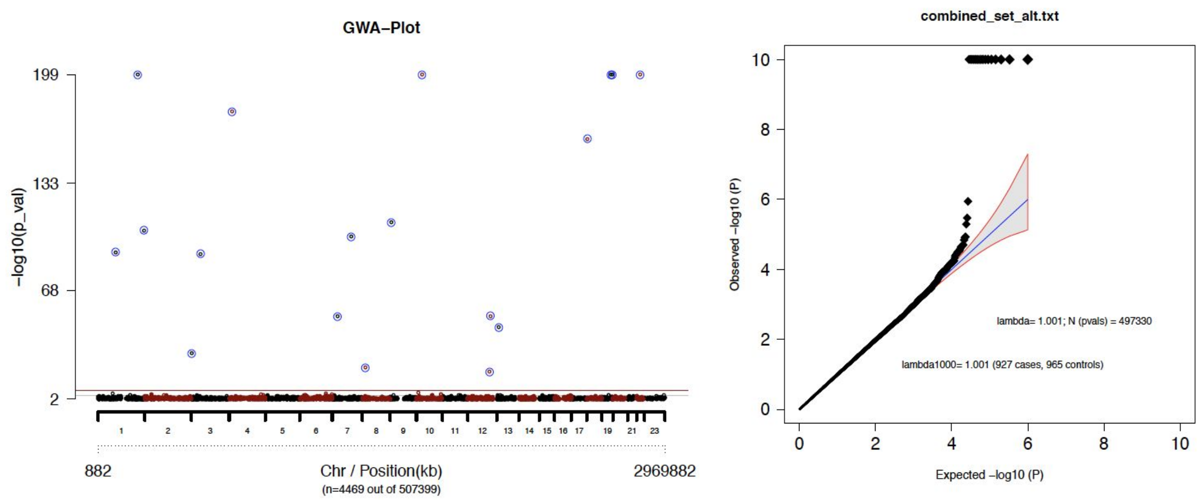


Extended Data Figure 9: Post-QC QQ plot from second *‘preimp’* on cohort 1

After removing the Asian subset from the cohort, we observed that the lambda inflation was reduced to an acceptable value (<1.05) (Note: Extended Data Figure 6 and Extended Data Figure 9). Nevertheless, there remained 28 genome-wide significant SNPs which required further quality ascertainment. A crucial feature of the RICOPILI *preimp* module includes GWAS summary statistics annotated with quality control parameters prior to imputation, which is provided in the **.detres* file. We were able to identify high missing rate for SNPs flagged as GWAS significant (Extended Data Figure 9). The following table (Extended Data Figure 10) was extracted using the command below.

```
-bash:~> awk '$8<5.0e-08 || NR ==1'
sim_hap1b_eur_sa-qc2.detres > sim_hap1b_eur_sa-qc2.detres.gws
-bash:~> column -t sim_hap1b_eur_sa-qc2.detres.gws
```

| RSID | CHR | BP | A1 | F_A | F_U | A2 | P_ASSOC | OR | SE | MAF | F_MISS | F_MISS_A | F_MISS_U | F_MISS_DIFF | F_MISS_P | log(P)_HWE_cas | log(P)_HWE_con | log(P)_HWE_all |
|---|---|---|---|---|---|---|---|---|---|---|---|---|---|---|---|---|---|---|
| rs10862292 | 12 | 81711953 | G | 0.9277 | 0.06955 | A | 0 | 171.7 | 0.1277 | 0.4903 | 0.01586 | 0.0151 | 0.01658 | -0.00148 | 0.8553 | 0 | -0.0967 | -246 |
| rs7611432 | 3 | 53971360 | G | 0.9571 | 0.0385 | T | 0 | 557.7 | 0.1663 | 0.4884 | 0.01797 | 0.01834 | 0.01762 | 0.000719999999999998 | 1 | -0.394 | 0 | -200 |
| rs4848495 | 2 | 118928678 | C | 0.9091 | 0.08281 | A | 0 | 110.8 | 0.1165 | 0.4882 | 0.01638 | 0.0151 | 0.01762 | -0.00252 | 0.7196 | -0.162 | -0.875 | -211 |
| rs221795 | 7 | 100283261 | C | 0.6663 | 0.3158 | T | 1.797e-101 | 4.326 | 0.07003 | 0.4874 | 0.01638 | 0.01726 | 0.01554 | 0.00172 | 0.8569 | -0.299 | -0.301 | -6.92 |
| rs6940091 | 6 | 2437996 | G | 0.8615 | 0.1452 | A | 0 | 36.63 | 0.09413 | 0.4962 | 0.0185 | 0.01834 | 0.01865 | -0.000310000000000001 | 1 | -0.313 | 0 | -115 |
| rs37267 | 7 | 41337044 | A | 0.8571 | 0.1126 | G | 0 | 47.3 | 0.09888 | 0.4776 | 0.01903 | 0.01834 | 0.01969 | -0.00135 | 0.8676 | -0.0495 | 0 | -133 |
| rs7524120 | 1 | 92865160 | G | 0.6559 | 0.3224 | A | 7.507e-92 | 4.006 | 0.06965 | 0.486 | 0.0185 | 0.01726 | 0.01969 | -0.00243 | 0.7353 | -0.517 | -0.0844 | -7.73 |
| rs11758794 | 6 | 161521680 | G | 0.9552 | 0.04198 | A | 0 | 487.1 | 0.1612 | 0.4925 | 0.0185 | 0.01187 | 0.02487 | -0.013 | 0.04052 | 0 | -0.395 | -200 |
| rs1417731 | 10 | 93635541 | G | 0.9071 | 0.08078 | A | 0 | 111.2 | 0.1168 | 0.4857 | 0.0185 | 0.01834 | 0.01865 | -0.000310000000000001 | 1 | 0 | -0.294 | -203 |
| rs7043262 | 9 | 6926738 | G | 0.676 | 0.3103 | A | 6.387e-110 | 4.638 | 0.07058 | 0.4889 | 0.01956 | 0.02265 | 0.01658 | 0.00607 | 0.4071 | -0.0271 | -0.705 | -10 |
| rs238112 | 18 | 3499767 | A | 0.7233 | 0.2802 | G | 8.939e-161 | 6.715 | 0.07318 | 0.4968 | 0.01691 | 0.01942 | 0.01451 | 0.00491 | 0.4771 | -0.168 | -0.0594 | -16.9 |
| rs1929254 | 10 | 30186233 | T | 0.7555 | 0.2466 | C | 3.847e-211 | 9.441 | 0.07626 | 0.4965 | 0.0185 | 0.01618 | 0.02073 | -0.00455 | 0.4986 | -0.184 | 0 | -29.3 |
| rs11234161 | 11 | 84469196 | T | 0.8812 | 0.1183 | G | 0 | 55.29 | 0.1016 | 0.4919 | 0.01903 | 0.01942 | 0.01865 | 0.00077 | 1 | 0 | -0.0576 | -144 |
| rs12533618 | 7 | 52142647 | A | 0.9078 | 0.1111 | G | 0 | 78.8 | 0.1091 | 0.4992 | 0.01691 | 0.02265 | 0.0114 | 0.01125 | 0.07353 | -0.161 | -1.67 | -184 |
| rs696964 | 1 | 208598450 | G | 0.7563 | 0.2282 | A | 1.04e-227 | 10.5 | 0.07727 | 0.4863 | 0.01691 | 0.01942 | 0.01451 | 0.00491 | 0.4771 | -0.492 | -0.706 | -26.6 |
| rs16947913 | 16 | 78554218 | C | 0.8069 | 0.1837 | A | 0 | 18.57 | 0.08391 | 0.4884 | 0.01744 | 0.01942 | 0.01554 | 0.00388 | 0.5995 | -0.47 | -0.0391 | -59.6 |
| rs149150 | 3 | 3094216 | G | 0.5857 | 0.3976 | A | 1.954e-30 | 2.142 | 0.06685 | 0.4898 | 0.0185 | 0.01834 | 0.01865 | -0.000310000000000001 | 1 | -0.077 | -1.04 | -0.215 |
| rs11715915 | 3 | 49455330 | T | 0.6625 | 0.3309 | C | 6.33e-91 | 3.969 | 0.06953 | 0.4933 | 0.01691 | 0.01726 | 0.01658 | 0.00068 | 1 | -0.427 | -0.146 | -6.37 |
| rs2157216 | 22 | 35344317 | A | 0.7968 | 0.1828 | G | 5.267e-308 | 17.54 | 0.08306 | 0.4834 | 0.01427 | 0.0151 | 0.01347 | 0.00163 | 0.8473 | -0.0368 | 0 | -60.6 |
| rs1248060 | 12 | 114864252 | C | 0.5667 | 0.4195 | T | 2.856e-19 | 1.81 | 0.06634 | 0.4919 | 0.01744 | 0.01294 | 0.02176 | -0.00882 | 0.1619 | -0.697 | -0.0487 | -0.0333 |
| rs4698491 | 4 | 16526736 | A | 0.7243 | 0.259 | G | 3.17e-177 | 7.516 | 0.07412 | 0.4876 | 0.01691 | 0.01402 | 0.01969 | -0.00567 | 0.3761 | -0.669 | -0.303 | -21.3 |
| rs11772815 | 7 | 28391047 | A | 0.3713 | 0.6225 | G | 5.805e-53 | 0.3582 | 0.06777 | 0.4992 | 0.01691 | 0.0151 | 0.01865 | -0.00355 | 0.5959 | -0.447 | -0.62 | -0.676 |
| rs168474 | 8 | 15599587 | G | 0.5728 | 0.4155 | A | 8.632e-22 | 1.886 | 0.0664 | 0.4927 | 0.01638 | 0.01402 | 0.01865 | -0.00463 | 0.4723 | -0.164 | -0.457 | -0.287 |
| rs275581 | 19 | 48848274 | C | 0.2123 | 0.7712 | A | 4.63e-255 | 0.07997 | 0.0791 | 0.4987 | 0.0148 | 0.01942 | 0.01036 | 0.00906 | 0.1275 | -0.312 | -0.504 | -35.9 |
| rs558107 | 13 | 30172458 | G | 0.6175 | 0.3833 | A | 2.595e-46 | 2.597 | 0.06743 | 0.4984 | 0.01586 | 0.01294 | 0.01865 | -0.00571 | 0.3607 | -0.849 | -0.167 | -0.979 |
| rs740032 | 12 | 120264341 | G | 0.6203 | 0.3684 | A | 3.096e-53 | 2.8 | 0.06779 | 0.4911 | 0.01691 | 0.02265 | 0.0114 | 0.01125 | 0.07353 | -0.0517 | -0.576 | -1.11 |
| rs10412597 | 19 | 56473189 | G | 0.2168 | 0.7592 | T | 5.888e-240 | 0.08779 | 0.0782 | 0.4935 | 0.01691 | 0.01726 | 0.01658 | 0.00068 | 1 | -0.073 | 0 | -37.3 |
| rs12140273 | 1 | 241542880 | C | 0.6786 | 0.321 | T | 3.096e-105 | 4.465 | 0.0703 | 0.4962 | 0.0185 | 0.01834 | 0.01865 | -0.000310000000000001 | 1 | -1.02 | -0.341 | -8.7 |

Extended Data Figure 10: Cohort 1: List of GWAS hits in second *'preimp'* association analysis.

#### File: sim_hap1b_eur_sa-qc2.pdf

##### 4.2 Lambda-Plot to various variables (all post-QC)

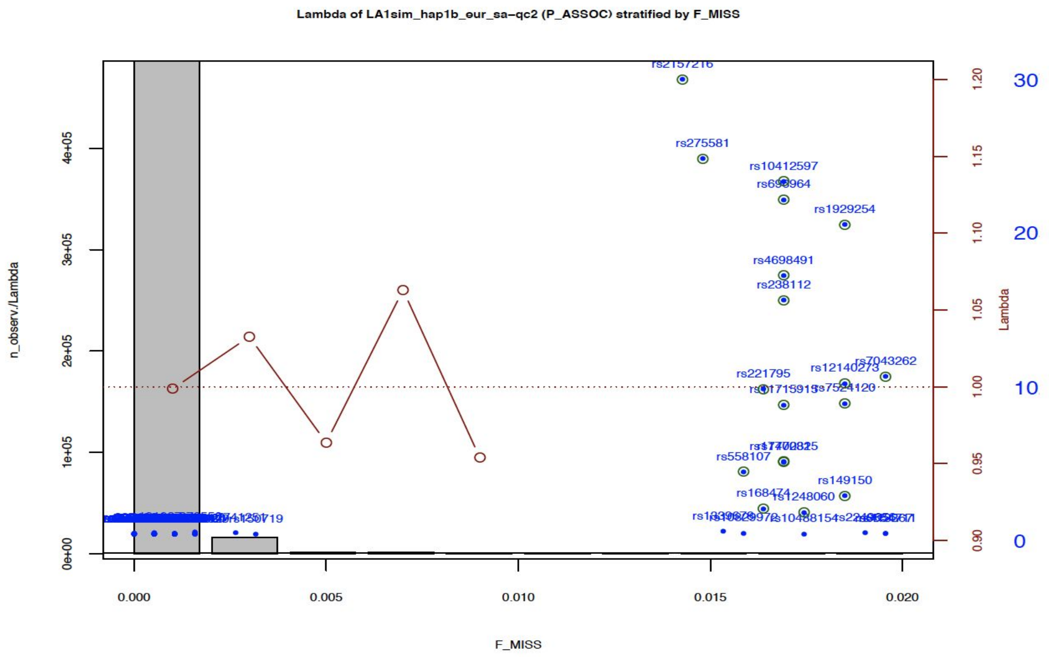


Extended Data Figure 11: Lambda stratified by missing rate.

RICOPILI generates the "lahunt" - plot (Extended Data Figure 11) to visualize various QC parameters (in this example: missing rate in SNPs) on the x-axis with regards to Lambda and genome-wide SNP -log10 p-values on the y-axes[40]: Grey bars (y-axis on the left) show the number of SNPs within binned values of the effect-variable. Lambda-GC within each bin was plotted with red line/dots (red right y-axis). The blue dots represent -log10(P) of the top associated SNPs (blue y-axis on the right).

[40] QC report includes by default: SNP missing rate (cases, controls, combined) and missing difference between cases and controls, minor allele frequency (MAF), Hardy-Weinberg equilibrium (cases, controls, combined), CHR.

Genome-wide significant signals might be present due to technical errors found within the data other than being a true genetic association. Being sufficiently confident that the GWAS signal was likely a false positive (e.g. small cohort size for a psychiatric trait), a threshold that would help exclude these variant was identified. In this example, significant GWAS signals were present in SNPs between 1-2% rates of missingness in either cases or controls. We applied a stricter threshold than the default value (1% instead of 2%) to exclude these SNPs using the following command:

```
-bash:~> awk '$13>.01 || $14>.01 {print $1}'
sim_hap1b_eur_sa-qc2.detres > snps.exclusion.list
```

Then, using plink --exclude to exclude 4,277 SNPs

```
-bash:~> plink --bfile sim_hap1b_eur_sa-qc2 --exclude
snps.exclusion.list --make-bed --out hap1b_snp.exc
```

We performed the third and final preimp analysis after removing the technical errors, population biases and false positives (hap1b_snp.exc.bed/.bim/.fam).

```
-bash:~> preimp_dir --dis sim --out hap1b_SNP.ex
```

After invoking the 'preimp' module, we updated the 'sim.names' with "hap1c" under the STUDYNAME column and "3" under QCCYCLE (since this was the third 'preimp' module on cohort 1; Extended Data Figure 12).

```
###STUDYNAME: 5 alphanumeric characters, recommeded city or study abbreviation and wave, e.g. lond1, cloz2
###               please negotiate with PIs about this name
###BFILE:     bed-filename root, e.g. if you dataset is called YOURDATA.bed, then please use YOURDATA in this column
###QCCYCLE:   numeric indicating the rounds of quality controls you have already performed
###               this will be appended to the resulting files with -qc1 or -qc2
###EXCLUDE:   set 0 if you want this dataset being included in the preimp process
STUDYNAME       BFILE           QCCYCLE         EXCLUDE
hap1c           hap1b_snp.exc     3               0
```

Extended Data Figure 12: Screenshot '*sim.names*' (*[--dis].names*) from third '*preimp*' on cohort 1.

We saved 'sim.names' ('[--dis].names') and repeated the 'preimp' command in the same directory.

```
-bash:~> preimp_dir --dis sim --out hap1b_SNP.ex
```

| S.No | Pre-Qc'd genotype file names | QC reports files names | Post-Qc'd genotype file names |
|---|---|---|---|
| 1 | hap1b_SNP.ex | sim_hap1c_eur_sa-qc3.pdf | sim_hap1c_eur_sa-qc3 |

Extended Data Table 4: Pre/Post QC files from third '*preimp*' on cohort 1

**File: sim_hap1c_eur_sa-qc3.pdf**

### 1 General Info

#### 1.1 Size of Sample

| Test | pre QC | post QC | exclusion-N |
|---|---|---|---|
| Cases,Controls,Missing | 927,965,0 | 927,965,0 | 0,0,0 |
| Males,Females,Unspec | 1166,726,0 | 1166,726,0 | 0,0,0 |
| SNPs | 503121 | 503121 | 0 (0.0%) |

#### 1.2 Exclusion overview

- would have excluded 0 individuals without pre-filter (SNP-Missing 0.05)

| Filter | N |
|---|---|
| SNPs: call rate < 0.950 (pre - filter) | 0 (0.0%) |
| IDs: call rate (cases/controls) < 0.980 | 0 (0/0) |
| IDs: FHET outside +- 0.20 (cases/controls) | 0 (0/0) |
| IDs: Sex violations -excluded- (N-tested) | 0 (1892) |
| IDs: Sex warnings (undefined phenotype / ambiguous genotypes) | 1 (0/1) |
| SNPs: call rate < 0.980 | 0 (0.0%) |
| SNPs: missing difference > 0.020 | 0 (0.0%) |
| SNPs: without valid association p-value (invariant) | 0 (0.0%) |
| SNPs: HWE-controls < 1e-06 | 0 (0.0%) |
| SNPs: HWE-cases < 1e-10 | 0 (0.0%) |
| Warning: genomewide significant SNPs (autosomal/known) | 0 (0/0) |

Extended Data Figure 13: Detailed QC report of third '*preimp*' on cohort 1

##### 2.2 Manhattan-Plot - post-QC (QQplot with MAF 0.02)

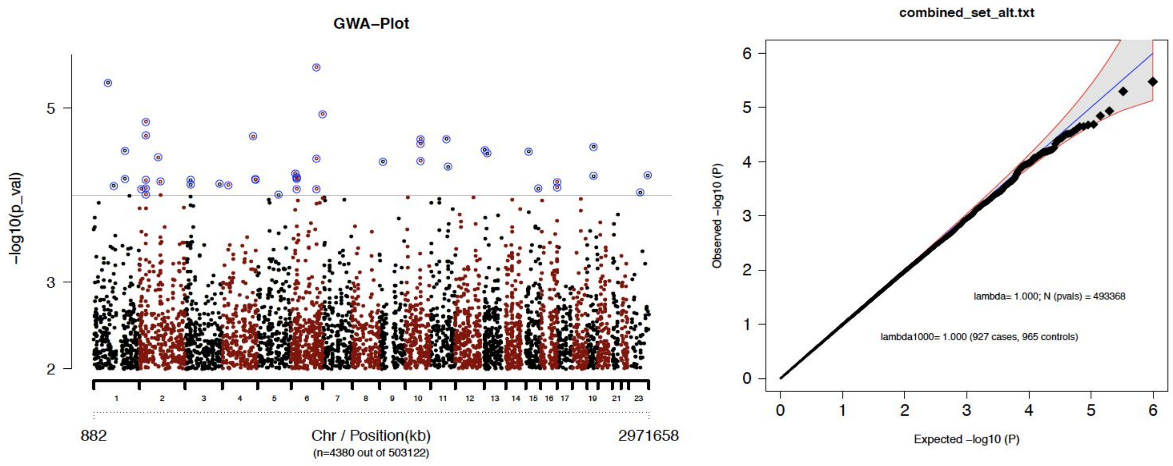


Extended Data Figure 14: Post-QC QQ plot from third *'preimp'* on cohort 1

Data reported in Extended Data Figures 13 and 14 demonstrate that with various QC actions we were able to achieve a quality controlled genotype dataset with no noticeable biases.

#### Cohort 2 (sim_sim2a_eur_sa_merge.miss.bed/.bim/.fam)

The second cohort consisted of 474 cases and 526 controls of European ancestry. In this dataset, we introduced technical errors described in section 2.

Preimp/QC results:

**File: sim_hap2a_eur_sa-qc1.pdf**

### 1 General Info

#### 1.1 Size of Sample

| Test | pre QC | post QC | exclusion-N |
|---|---|---|---|
| Cases,Controls,Missing | 474,526,0 | 423,477,0 | 51,49,0 |
| Males,Females,Unspec | 605,395,0 | 548,352,0 | 57,43,0 |
| SNPs | 593970 | 556348 | 37622 (6.3%) |

#### 1.2 Exclusion overview

- would have excluded 67 individuals without pre-filter (SNP-Missing 0.05)

| Filter | N |
|---|---|
| SNPs: call rate < 0.950 (pre - filter) | 138 (0.0%) |
| IDs: call rate (cases/controls) < 0.980 | 67 (34/33) |
| IDs: FHET outside +- 0.20 (cases/controls) | 20 (10/10) |
| IDs: Sex violations -excluded- (N-tested) | 28 (1000) |
| IDs: Sex warnings (undefined phenotype / ambiguous genotypes) | 0 (0/0) |
| SNPs: call rate < 0.980 | 8433 (1.4%) |
| SNPs: missing difference > 0.020 | 6181 (1.1%) |
| SNPs: without valid association p-value (invariant) | 1 (0.0%) |
| SNPs: HWE-controls < -6 | 11676 (2.0%) |
| SNPs: HWE-cases < -10 | 11677 (2.0%) |
| Warning: genomewide significant SNPs (autosomal/known) | 29 (29/0) |

Extended Data Figure 15: Detailed QC report of first *'preimp'* on cohort 2

#### 2 Manhattan

##### 2.1 Manhattan-Plot - pre-QC (QQplot with MAF 0.02)

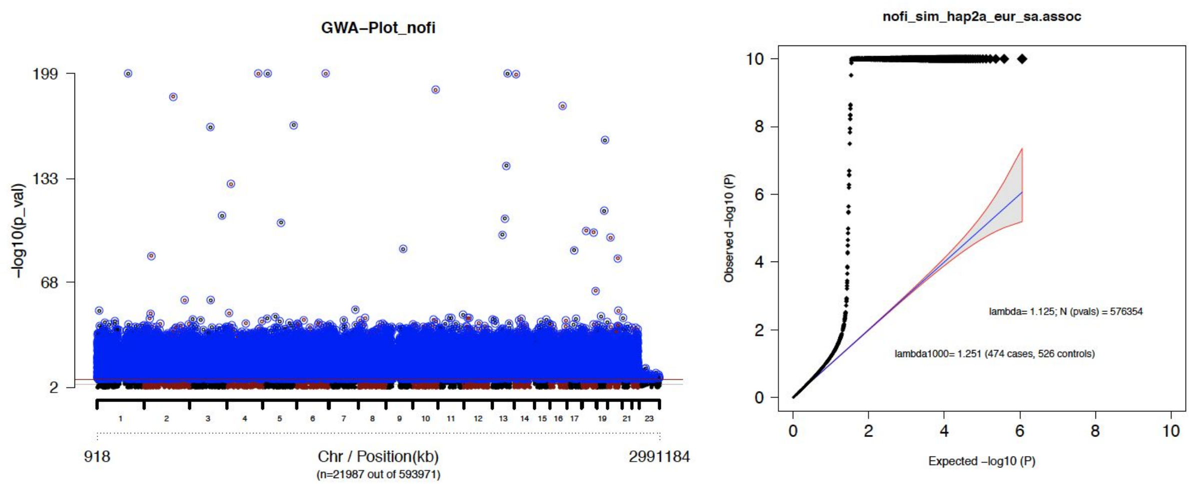


Extended Data Figure 16:: Pre-QC QQ plot from first *'preimp'* on cohort 2.

##### 2.2 Manhattan-Plot - post-QC (QQplot with MAF 0.02)

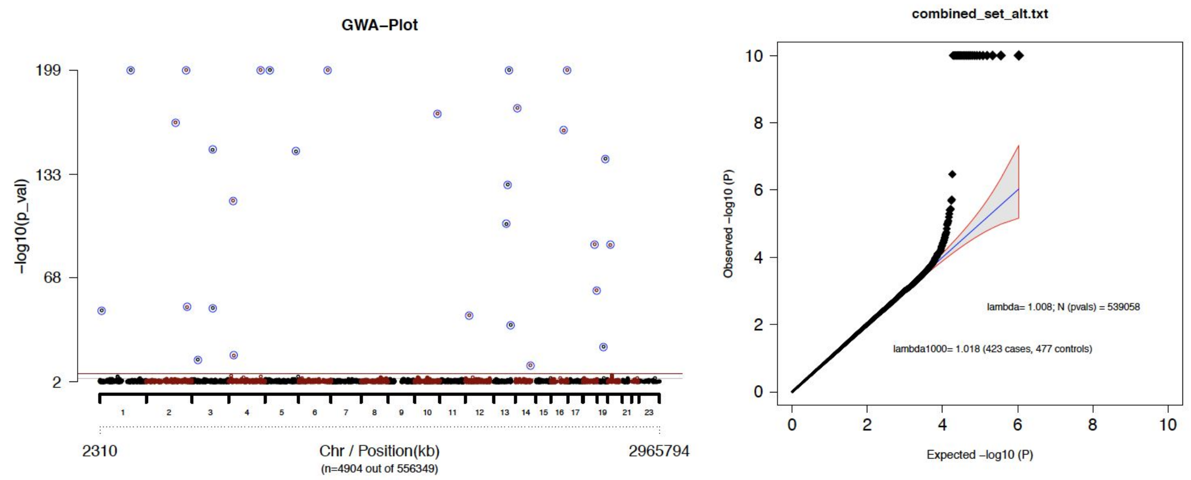


Extended Data Figure 17: Post-QC QQ plot from first *'preimp'* on cohort 2.

The default setting of the 'preimp' module excluded 37,622 (6.3%) of SNPs and 100 individuals, for technical errors (Extended Data Figure 15). We noted that Lambda inflation was resolved (Extended Data Figures 16 and 17) from default mode quality control. There were 29 genome-wide significant SNPs in the QC report, which required further quality control ascertainment.

```
-bash:~> awk '$8<5.0e-08 || NR ==1'
sim_hap2a_eur_sa-qc1.detres > sim_hap2a_eur_sa-qc1.detres.gws
column -t sim_hap2a_eur_sa-qc1.detres.gws
```

| RSID | CHR | BP | A1 | F_A | F_U | A2 | P_ASSOC | OR | SE | MAF | F_MISS | F_MISS_A | F_MISS_U | F_MISS_DIFF | F_MISS_P | log(P)_HWE_cas | log(P)_HWE_con | log(P)_HWE_all |
|---|---|---|---|---|---|---|---|---|---|---|---|---|---|---|---|---|---|---|
| rs9818848 | 3 | 112394340 | T | 0.3125 | 0.6635 | C | 2.876e-49 | 0.2305 | 0.1017 | 0.4989 | 0.01444 | 0.01655 | 0.01258 | 0.00397 | 0.7811 | -0.513 | -0.27 | -3.78 |
| rs3743739 | 16 | 68293320 | C | 0.1558 | 0.8024 | T | 8.746e-162 | 0.04546 | 0.1262 | 0.4989 | 0.02 | 0.02128 | 0.01887 | 0.00241 | 0.816 | 0 | -0.61 | -32.1 |
| rs7143061 | 14 | 100100164 | G | 0.4065 | 0.5798 | A | 3.176e-13 | 0.4964 | 0.09663 | 0.4983 | 0.01444 | 0.01418 | 0.01468 | -0.0005 | 1 | -0.651 | -0.152 | -1.29 |
| rs1581089 | 4 | 26155360 | C | 0.6024 | 0.3872 | T | 1.599e-19 | 2.398 | 0.09754 | 0.4881 | 0.01667 | 0.01891 | 0.01468 | 0.00423 | 0.7953 | -0.0368 | -1.19 | -2.15 |
| rs4622605 | 18 | 73139008 | C | 0.6847 | 0.2947 | T | 1.578e-60 | 5.197 | 0.1033 | 0.478 | 0.01444 | 0.01418 | 0.01468 | -0.0005 | 1 | -0.511 | -0.0401 | -6.22 |
| rs6793913 | 3 | 33029986 | T | 0.5778 | 0.3801 | C | 9.172e-17 | 2.232 | 0.0972 | 0.4734 | 0.01667 | 0.01182 | 0.02096 | -0.00914 | 0.3109 | -1.91 | -0.255 | -1.74 |
| rs9524001 | 13 | 93947095 | A | 0.7837 | 0.212 | G | 2.775e-127 | 13.46 | 0.1162 | 0.4813 | 0.01889 | 0.01655 | 0.02096 | -0.00441 | 0.8072 | -1.22 | -0.169 | -24.6 |
| rs3746319 | 19 | 44612231 | G | 0.8031 | 0.1951 | A | 1.102e-143 | 16.83 | 0.1201 | 0.4802 | 0.01889 | 0.02128 | 0.01677 | 0.00451 | 0.6336 | -0.193 | -0.623 | -29.1 |
| rs704193 | 12 | 22015768 | G | 0.6622 | 0.328 | A | 1.034e-44 | 4.016 | 0.1011 | 0.4842 | 0.01778 | 0.02364 | 0.01258 | 0.01106 | 0.3122 | -0.18 | -0.122 | -3.02 |
| rs11001589 | 10 | 77782036 | C | 0.9505 | 0.03715 | A | 0 | 497.4 | 0.2352 | 0.4644 | 0.01667 | 0.02128 | 0.01258 | 0.0087 | 0.4351 | -0.212 | -0.318 | -156 |
| rs4491070 | 1 | 12130051 | T | 0.6735 | 0.3277 | C | 8.463e-48 | 4.233 | 0.1015 | 0.4898 | 0.01667 | 0.01891 | 0.01468 | 0.00423 | 0.7953 | -0.0404 | 0 | -3.44 |
| rs28608705 | 22 | 43897310 | G | 0.9758 | 0.01589 | A | 0 | 2502 | 0.3449 | 0.4644 | 0.01556 | 0.02128 | 0.01048 | 0.0108 | 0.2804 | 0 | 0 | -203 |
| rs10886725 | 10 | 122386059 | A | 0.8481 | 0.1827 | G | 4.231e-172 | 24.98 | 0.1282 | 0.4966 | 0.01556 | 0.01182 | 0.01887 | -0.00705 | 0.4322 | -0.462 | -0.121 | -40.6 |
| rs16896876 | 5 | 27523340 | G | 0.9118 | 0.07569 | T | 1.489e-270 | 126.3 | 0.174 | 0.4677 | 0.01889 | 0.02128 | 0.01677 | 0.00451 | 0.6336 | -1.54 | -1.41 | -115 |
| rs7536863 | 1 | 165953874 | G | 0.8675 | 0.1002 | A | 6.456e-229 | 58.77 | 0.1493 | 0.4604 | 0.01778 | 0.01891 | 0.01677 | 0.00214 | 0.8072 | -0.285 | -0.218 | -72.3 |
| rs8098946 | 18 | 62506122 | G | 0.7356 | 0.258 | A | 1.248e-89 | 8.002 | 0.1084 | 0.4836 | 0.01667 | 0.01182 | 0.02096 | -0.00914 | 0.3109 | -0.0954 | -0.0439 | -10 |
| rs1963996 | 4 | 22715639 | G | 0.7669 | 0.2197 | A | 4.741e-117 | 11.68 | 0.1138 | 0.4757 | 0.01667 | 0.02128 | 0.01258 | 0.0087 | 0.4351 | -1.71 | -0.162 | -13.3 |
| rs7259736 | 19 | 35096264 | G | 0.6286 | 0.3838 | A | 6.646e-25 | 2.717 | 0.09802 | 0.4994 | 0.01222 | 0.007092 | 0.01677 | -0.009678 | 0.233 | -0.333 | -0.114 | -0.459 |
| rs10457902 | 6 | 154297720 | G | 0.8627 | 0.1375 | T | 8.199e-204 | 39.39 | 0.1384 | 0.4779 | 0.01778 | 0.01891 | 0.01677 | 0.00214 | 0.8072 | -0.677 | -0.363 | -62.3 |
| rs1601527 | 2 | 156261310 | A | 0.8329 | 0.178 | C | 1.382e-166 | 23.03 | 0.1263 | 0.4836 | 0.01667 | 0.02364 | 0.01048 | 0.01316 | 0.1907 | -0.319 | -0.195 | -33.9 |
| rs9865917 | 3 | 111167008 | A | 0.799 | 0.1794 | G | 1.413e-149 | 18.19 | 0.1214 | 0.4689 | 0.01778 | 0.02364 | 0.01258 | 0.01106 | 0.3122 | -0.66 | -0.273 | -34.6 |
| rs7986007 | 13 | 100702118 | G | 0.8957 | 0.113 | A | 1.435e-237 | 67.39 | 0.1532 | 0.4814 | 0.01556 | 0.01418 | 0.01677 | -0.00259 | 0.7942 | -0.101 | -0.0868 | -77.7 |
| rs9312517 | 4 | 168139380 | G | 0.8978 | 0.1156 | A | 2.001e-236 | 67.21 | 0.1535 | 0.4841 | 0.01889 | 0.01655 | 0.02096 | -0.00441 | 0.8072 | -0.985 | -0.307 | -71.2 |
| rs17413177 | 2 | 212419760 | T | 0.9531 | 0.06103 | C | 5.389e-307 | 312.8 | 0.2135 | 0.4813 | 0.01889 | 0.01655 | 0.02096 | -0.00441 | 0.8072 | -0.217 | -0.396 | -137 |
| rs2327886 | 20 | 14724600 | T | 0.744 | 0.266 | C | 2.499e-89 | 8.017 | 0.1087 | 0.4904 | 0.02 | 0.02128 | 0.01887 | 0.00241 | 0.816 | -1.7 | -0.622 | -9.28 |
| rs10146422 | 14 | 30839557 | C | 0.8301 | 0.1578 | T | 1.156e-175 | 26.08 | 0.1287 | 0.4734 | 0.01778 | 0.01891 | 0.01677 | 0.00214 | 0.8072 | -0.22 | -0.642 | -42.7 |
| rs10851225 | 13 | 109659839 | A | 0.6574 | 0.3468 | G | 8.703e-39 | 3.614 | 0.1004 | 0.4921 | 0.01889 | 0.02364 | 0.01468 | 0.00896 | 0.3393 | -0.351 | -0.813 | -1.66 |
| rs284544 | 2 | 217309111 | A | 0.6867 | 0.3319 | G | 4.618e-50 | 4.413 | 0.1021 | 0.4989 | 0.02 | 0.01891 | 0.02096 | -0.00205 | 1 | 0 | -0.122 | -4.15 |
| rs3922878 | 16 | 86854704 | A | 0.9457 | 0.05745 | C | 6.531e-305 | 285.5 | 0.2077 | 0.4734 | 0.01778 | 0.02128 | 0.01468 | 0.0066 | 0.4632 | -0.469 | -0.411 | -136 |
| rs10475761 | 5 | 165309208 | A | 0.8022 | 0.1848 | G | 1.425e-148 | 17.88 | 0.121 | 0.4757 | 0.01667 | 0.01418 | 0.01887 | -0.00469 | 0.6141 | -0.0569 | -0.901 | -33.7 |
| rs675350 | 13 | 86327523 | A | 0.7482 | 0.2367 | G | 1.223e-102 | 9.58 | 0.1108 | 0.4763 | 0.01556 | 0.01891 | 0.01258 | 0.00633 | 0.5913 | 0 | -0.794 | -16.9 |

Extended Data Figure 18: Cohort 2: List of GWAS hits in second *'preimp'* association analysis

The report in Extended Data Figure 15 revealed SNPs with high missing rate in significant variants. We applied a stricter filter threshold that removed 4,451 additional SNPs from the cohort on post QC'd genotypes cohort 2 (sim_hap2a_eur_sa-qc1.bed/.bim/.fam; Extended Data Table 2).

```
-bash:~> awk '$13>.01 || $14>.01 {print $1}'
sim_hap2a_eur_sa-qc1.detres > snps.exclusion.list
```

```
-bash:~> plink --bfile sim_hap2a_eur_sa-qc1 --exclude
snps.exclusion.list --make-bed --out hap2a_snp.exc
```

The second 'preimp' on the filtered genotype data with the false positive SNPs (hap2a_snp.exc.bed/.bim/.fam) was carried out.

```
-bash: preimp_dir --dis sim --out hap2a_SNP.ex
```

```
###STUDYNAME: 5 alphanumeric characters, recommeded city or study abbreviation and wave, e.g. lond1, cloz2
###              please negotiate with PIs about this name
###BFILE:      bed-filename root, e.g. if you dataset is called YOURDATA.bed, then please use YOURDATA in this column
###QCCYCLE:    numeric indicating the rounds of quality controls you have already performed
###              this will be appended to the resulting files with -qc1 or -qc2
###EXCLUDE:    set 0 if you want this dataset being included in the preimp process
STUDYNAME            BFILE         QCCYCLE       EXCLUDE
hap2b            hap2a_snp.exc        2             0
```

Extended Data Figure 19: Screenshot '*sim.names*' (*[--dis].names*) from second '*preimp*' on cohort 2.

We modified 'sim.names' ('[--dis].names') with details found in  Extended Data Figure 19 and repeated the 'preimp' command in the same directory.

```
-bash: preimp_dir --dis sim --out hap2a_SNP.ex
```

| S.No | Pre-Qc'd genotype file names | QC reports files names | Post-Qc'd genotype file names |
|---|---|---|---|
| 1 | hap2a_snp.exc | sim_hap2b_eur_sa-qc2.pdf | sim_hap2b_eur_sa-qc2 |

Extended Data Table 5: Pre/Post QC files from the second '*preimp*' on cohort 2

**File: sim_hap2b_eur_sa-qc2.pdf**

### 1 General Info

#### 1.1 Size of Sample

| Test | pre QC | post QC | exclusion-N |
|---|---|---|---|
| Cases,Controls,Missing | 423,477,0 | 423,477,0 | 0,0,0 |
| Males,Females,Unspec | 548,352,0 | 548,352,0 | 0,0,0 |
| SNPs | 551898 | 551898 | 0 (0.0%) |

#### 1.2 Exclusion overview

- would have excluded 0 individuals without pre-filter (SNP-Missing 0.05)

| Filter | N |
|---|---|
| SNPs: call rate < 0.950 (pre - filter) | 0 (0.0%) |
| IDs: call rate (cases/controls) < 0.980 | 0 (0/0) |
| IDs: FHET outside +- 0.20 (cases/controls) | 0 (0/0) |
| IDs: Sex violations -excluded- (N-tested) | 0 (900) |
| IDs: Sex warnings (undefined phenotype / ambiguous genotypes) | 0 (0/0) |
| SNPs: call rate < 0.980 | 0 (0.0%) |
| SNPs: missing difference > 0.020 | 0 (0.0%) |
| SNPs: without valid association p-value (invariant) | 0 (0.0%) |
| SNPs: HWE-controls < -6 | 0 (0.0%) |
| SNPs: HWE-cases < -10 | 0 (0.0%) |
| Warning: genomewide significant SNPs (autosomal/known) | 0 (0/0) |

Extended Data Figure 20: Detailed QC report of second *'preimp'* on cohort 2

##### 2.2 Manhattan-Plot - post-QC (QQplot with MAF 0.02)

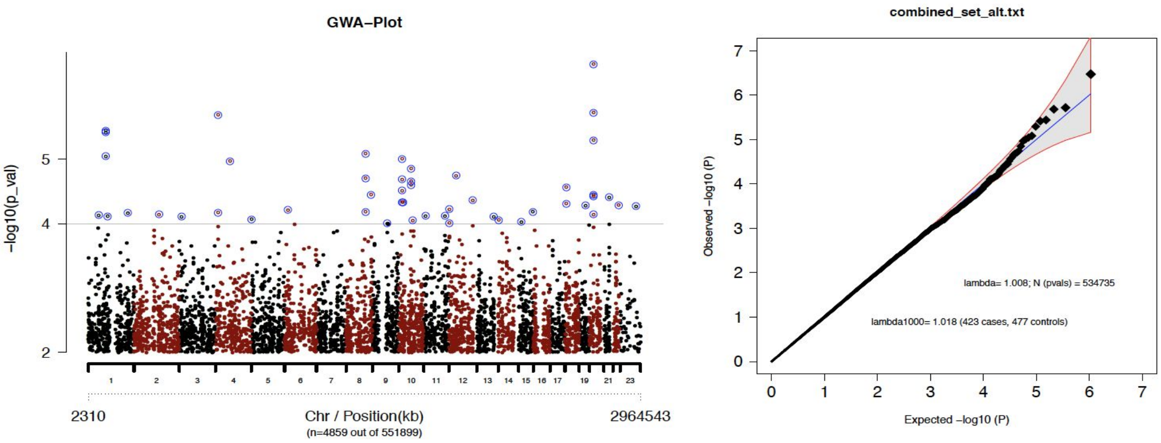


Extended Data Figure 21: Post-QC QQ plot from second *'preimp'* on cohort 2.

After the second 'preimp' quality control procedure for cohort 2 (Extended Figures 20 and 21 a reasonably QC-ed dataset without spurious results SNPs (551,898) was available for imputation.

#### Cohort 3 (hapgen_sample3a.bed/.bim/.fam)

Cohort 3 consists of 473 cases and 527 controls of European ancestry and 5 cases and 95 controls with Asian ancestry.

#### Preimp/QC results:

**File: sim_hap3a_eur_sa-qc1.pdf**

### 1 General Info

#### 1.1 Size of Sample

| Test | pre QC | post QC | exclusion-N |
|---|---|---|---|
| Cases,Controls,Missing | 478,622,0 | 478,622,0 | 0,0,0 |
| Males,Females,Unspec | 633,467,0 | 633,467,0 | 0,0,0 |
| SNPs | 547764 | 540914 | 6850 (1.3%) |

#### 1.2 Exclusion overview

- would have excluded 0 individuals without pre-filter (SNP-Missing 0.05)

| Filter | N |
|---|---|
| SNPs: call rate < 0.950 (pre - filter) | 0 (0.0%) |
| IDs: call rate (cases/controls) < 0.980 | 0 (0/0) |
| IDs: FHET outside +- 0.20 (cases/controls) | 0 (0/0) |
| IDs: Sex violations -excluded- (N-tested) | 0 (1100) |
| IDs: Sex warnings (undefined phenotype / ambiguous genotypes) | 0 (0/0) |
| SNPs: call rate < 0.980 | 0 (0.0%) |
| SNPs: missing difference > 0.020 | 0 (0.0%) |
| SNPs: without valid association p-value (invariant) | 0 (0.0%) |
| SNPs: HWE-controls < 1e-06 | 6850 (1.3%) |
| SNPs: HWE-cases < 1e-10 | 0 (0.0%) |
| Warning: genomewide significant SNPs (autosomal/known) | 0 (0/0) |

Extended Data Figure 22: Detailed QC report of first *'preimp'* on cohort 3

#### 2 Manhattan

##### 2.1 Manhattan-Plot - pre-QC (QQplot with MAF 0.02)

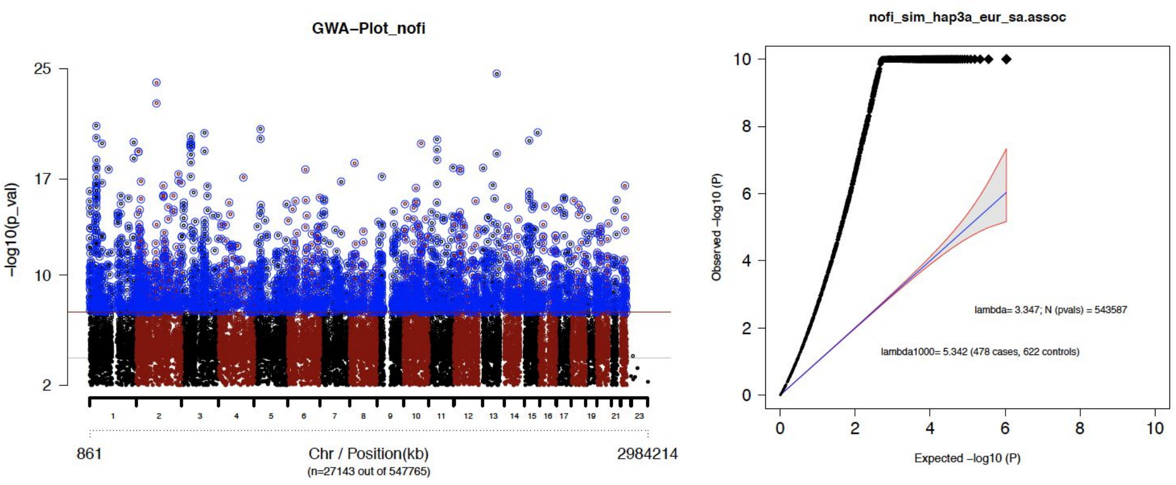


Extended Data Figure 23: Pre-QC QQ plot from first *'preimp'* on cohort 3.

##### 2.2 Manhattan-Plot - post-QC (QQplot with MAF 0.02)

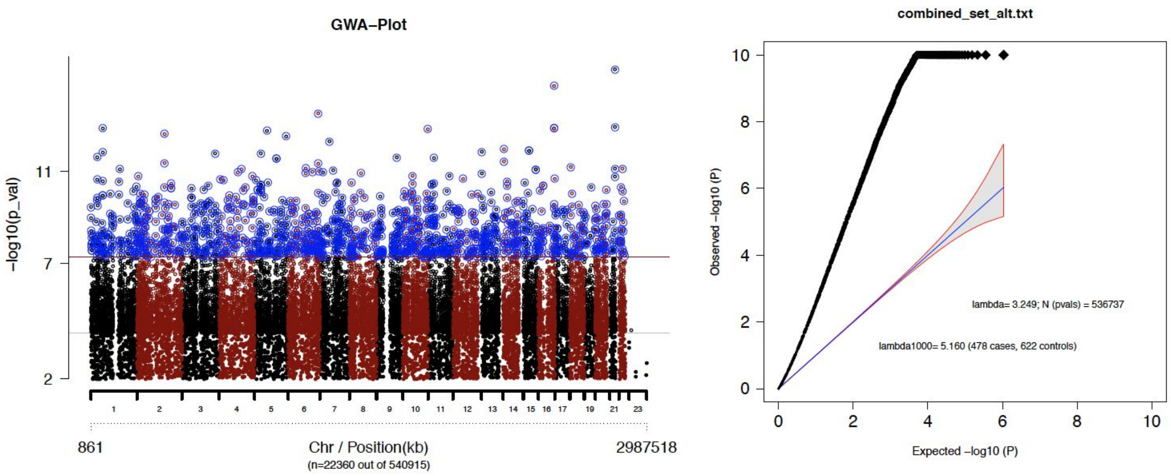


Extended Data Figure 24: Post-QC QQ plot from first *'preimp'* on cohort 3.

This QC report was relatively clean compared to cohort 1 and cohort 2. No individuals were excluded and a small fraction of SNPs (1.3 % - 6850) were excluded due to HWE - errors in controls (Extended Data Figure 22). No genome-wide significant SNPs were identified in the post-qc association analysis (Extended Data Figure 24). Nonetheless, it is notable that the QQ plot was highly inflated, suggesting the presence of population stratification errors related to the inclusion of the Asian individuals in this cohort. A PCA plot identifies individuals with European and Asian ancestries.

```
-bash:~> pcaer --prefercase --preferfam --out pca_hap3a
sim_hap3a_eur_sa-qc1.bim
```

#### Principal component analysis

**File pca_hap3a.menv.mds.2d.pdf.gz**

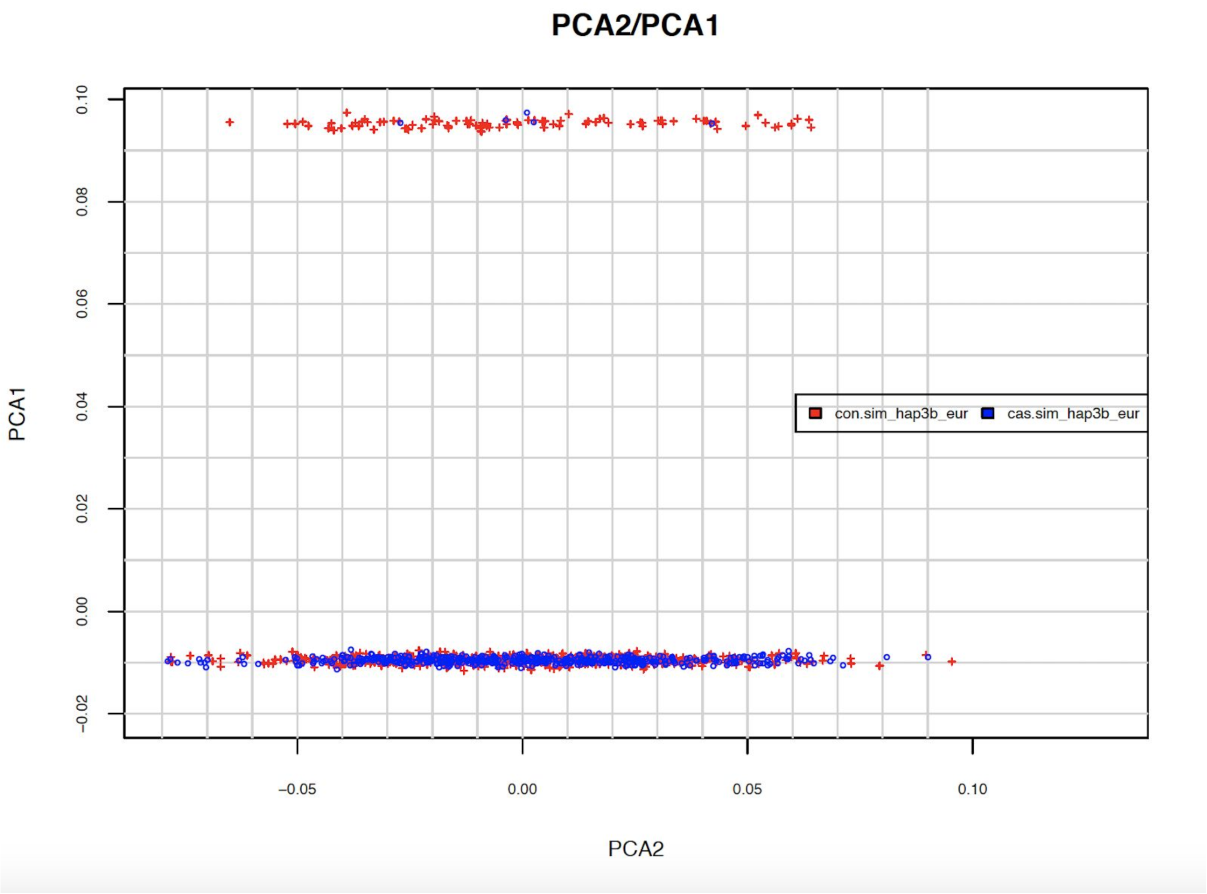


Extended Data Figure 25: PCA plot for cohort 3

The PCA plot (Extended Data Figure 25) shows two different ancestry clusters. For purpose of the current analysis we included individuals with European ancestry using a threshold on the first principal component i.e. PC-1 < 0.01 (use pca_hap3a.menv.mds). [41]

```
-bash:~> awk '$4<0.01' pca_hap3a.menv.mds >
hap3a.eur.sample.txt
```

These individuals were included using plink --keep. Thereafter, we performed another *'preimp'* procedure (Extended Data Figure 26, Extended Data Table 6) to check potential Lambda inflation (Extended Data Figure 27), which was reduced within normal range (1.006) after population stratification errors were resolved.

```
-bash:~> plink --bfile sim_hap3a_eur_sa-qc1 --keep
hap3a.eur.sample.txt --make-bed --out hap3a_only_eur
```

[41] RICOPILI leaves it to the user to decide which thresholds to use. On request we share scripts that do case-control matchings based on euclid distance.

```
-bash:~> preimp_dir --dis sim --out hap3a_only_eur
```

```
###STUDYNAME: 5 alphanumeric characters, recommeded city or study abbreviation and wave, e.g. lond1, cloz2
###              please negotiate with PIs about this name
###BFILE:      bed-filename root, e.g. if you dataset is called YOURDATA.bed, then please use YOURDATA in this column
###QCCYCLE:    numeric indicating the rounds of quality controls you have already performed
###              this will be appended to the resulting files with -qc1 or -qc2
###EXCLUDE:    set 0 if you want this dataset being included in the preimp process
STUDYNAME       BFILE         QCCYCLE     EXCLUDE
hap3b        hap3a_only_eur      2           0
```

Extended Data Figure 26: Screenshot '*sim.names*' (*[--dis].names*) from second '*preimp*' on cohort 3.

```
-bash preimp_dir --dis sim --out hap3a_only_eur
```

| S.No | Pre-Qc'd genotype file names | QC reports files names | Post-Qc'd genotype file names |
|---|---|---|---|
| 1 | hap3a_only_eur | sim_hap3b_eur_sa-qc2.pdf | sim_hap3b_eur_sa-qc2 |

Extended Data Table 6: Pre/Post QC files from second '*preimp*' on cohort 3

**File: sim_hap3b_eur_sa-qc2.pdf**

2.2 Manhattan-Plot - post-QC (QQplot with MAF 0.02)

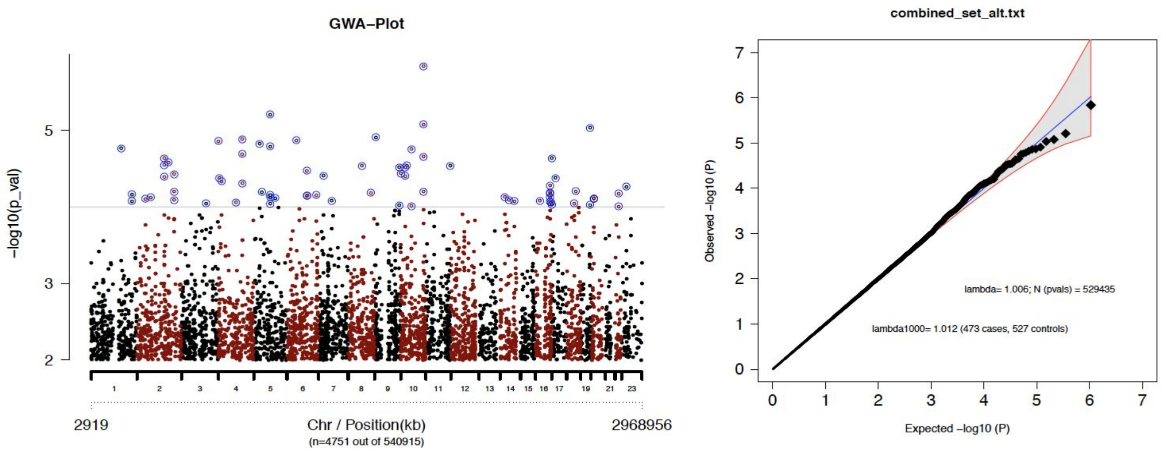


Extended Data Figure 27: Post-QC QQ plot from second *'preimp'* on cohort 3.

#### Cohort 4 (hapgen_sample4b.bed/.bim/fam) and Cohort 5 (hapgen_sample5a.ph.bed/.bim/fam)

We did not introduce technical or population biases to Cohorts 4 and 5. Cohort 4 comprised of 483 cases and 517 controls while cohort 5 contains 516 cases and 484 controls. Individuals in either cohorts were of European ancestry. Cohort 4 had 10 individuals overlapping with those in Cohort 5. We would demonstrate in this section how RICOPILI would identify sample overlaps across cohorts and the strategy we took to resolve these overlaps

Preimp/QC results:

**File sim_hap4a_eur_sa-qc1.pdf**

### 1 General Info

#### 1.1 Size of Sample

| Test | pre QC | post QC | exclusion-N |
|---|---|---|---|
| Cases,Controls,Missing | 483,517,0 | 483,517,0 | 0,0,0 |
| Males,Females,Unspec | 645,355,0 | 645,355,0 | 0,0,0 |
| SNPs | 593970 | 593970 | 0 (0.0%) |

#### 1.2 Exclusion overview

- would have excluded 0 individuals without pre-filter (SNP-Missing 0.05)

| Filter | N |
|---|---|
| SNPs: call rate < 0.950 (pre - filter) | 0 (0.0%) |
| IDs: call rate (cases/controls) < 0.980 | 0 (0/0) |
| IDs: FHET outside +- 0.20 (cases/controls) | 0 (0/0) |
| IDs: Sex violations -excluded- (N-tested) | 0 (1000) |
| IDs: Sex warnings (undefined phenotype / ambiguous genotypes) | 0 (0/0) |
| SNPs: call rate < 0.980 | 0 (0.0%) |
| SNPs: missing difference > 0.020 | 0 (0.0%) |
| SNPs: without valid association p-value (invariant) | 0 (0.0%) |
| SNPs: HWE-controls < 1e-06 | 0 (0.0%) |
| SNPs: HWE-cases < 1e-10 | 0 (0.0%) |
| Warning: genomewide significant SNPs (autosomal/known) | 0 (0/0) |

Extended Data Figure 28: Detailed QC report of first *'preimp'* on cohort 4

### 2 Manhattan

#### 2.1 Manhattan-Plot - pre-QC (QQplot with MAF 0.02)

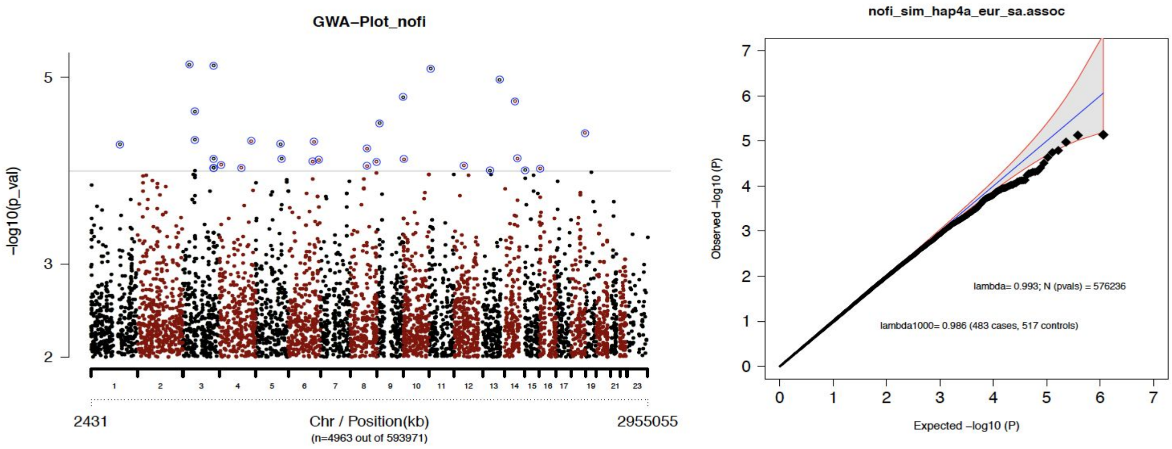


Extended Data Figure 29: Pre-QC QQ plot from first *'preimp'* on cohort 4.

**File sim_hap5a_eur_sa-qc1.pdf**

### 1 General Info

#### 1.1 Size of Sample

| Test | pre QC | post QC | exclusion-N |
|---|---|---|---|
| Cases,Controls,Missing | 516,484,0 | 516,484,0 | 0,0,0 |
| Males,Females,Unspec | 625,375,0 | 625,375,0 | 0,0,0 |
| SNPs | 593970 | 593970 | 0 (0.0%) |

#### 1.2 Exclusion overview

- would have excluded 0 individuals without pre-filter (SNP-Missing 0.05)

| Filter | N |
|---|---|
| SNPs: call rate < 0.950 (pre - filter) | 0 (0.0%) |
| IDs: call rate (cases/controls) < 0.980 | 0 (0/0) |
| IDs: FHET outside +- 0.20 (cases/controls) | 0 (0/0) |
| IDs: Sex violations -excluded- (N-tested) | 0 (1000) |
| IDs: Sex warnings (undefined phenotype / ambiguous genotypes) | 0 (0/0) |
| SNPs: call rate < 0.980 | 0 (0.0%) |
| SNPs: missing difference > 0.020 | 0 (0.0%) |
| SNPs: without valid association p-value (invariant) | 0 (0.0%) |
| SNPs: HWE-controls < -6 | 0 (0.0%) |
| SNPs: HWE-cases < -10 | 0 (0.0%) |
| Warning: genomewide significant SNPs (autosomal/known) | 0 (0/0) |

Extended Data Figure 30: Detailed QC report of first *'preimp'* on cohort 5.

### 2 Manhattan

#### 2.1 Manhattan-Plot - pre-QC (QQplot with MAF 0.02)

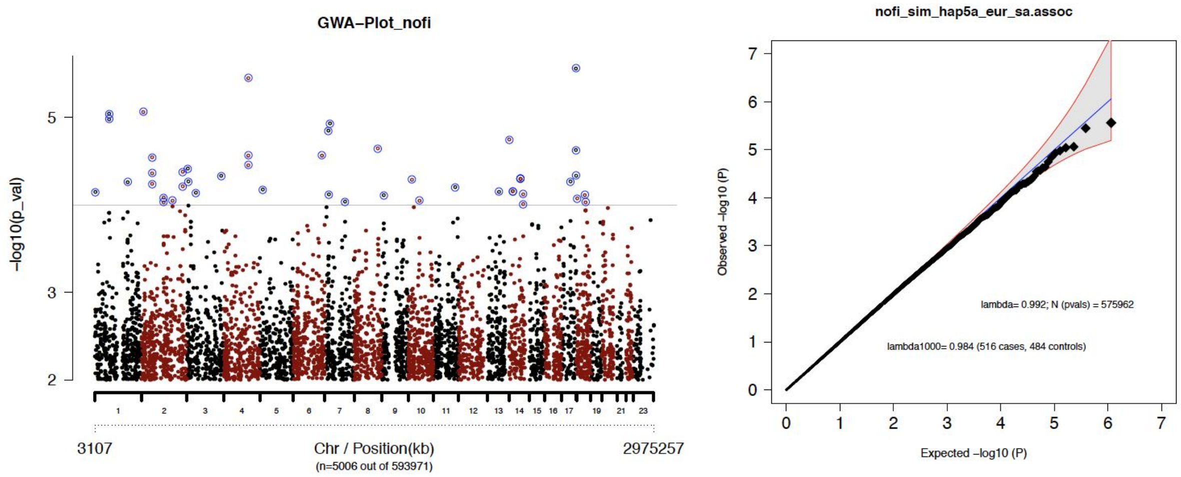


Extended Data Figure 31: Pre-QC QQ plot from first *'preimp'* on cohort 5.

QC reports generated by RICOPILI for cohorts 4 and 5 (Extended Data Figure 28; cohort 4 and Extended Data Figure 30; cohort 5) indicate that either data sets were relatively "clean". We expected these quality control outcomes since no interventions were introduced to these cohorts.

QQ plots for both cohorts are normal indicating the absence of population biases (Extended Data Figures 29 and 31). No further quality control procedures were required in these cohorts and was deemed appropriate for further analysis.

| QC Steps | Input Datasets | Data filtering | Output Datasets |
|---|---|---|---|
| 1 | sim_sim1a_eur_sa_merge.miss<br>sim_sim2a_eur_sa_merge.miss<br>hapgen_sample3a<br>hapgen_sample4b<br>hapgen_sample5a.ph | RICOPILI : *preimp* (Default parameters) | sim_hap1a_eur_sa-qc1<br>sim_hap2a_eur_sa-qc1<br>sim_hap3a_eur_sa-qc1<br>sim_hap4a_eur_sa-qc1*<br>sim_hap5a_eur_sa-qc1*<br>Note: eur = EUR population<br>sa = user initials |
| 2 | Cohort 1 (Second QC)<br>sim_hap1a_eur_sa-qc1 | Plink --keep | hap1a_only_eur |
| 3 | Cohort 1 (Second QC)<br>hap1a_only_eur | RICOPILI : *preimp* (Default parameters) | sim_hap1b_eur_sa-qc2 |
| 4 | Cohort 1 (Third QC)<br>sim_hap1b_eur_sa-qc2 | Plink --exclude | hap1b_SNP.ex |
| 5 | Cohort 1 (Third QC)<br>hap1b_SNP.ex | RICOPILI : *preimp* (Default parameters) | sim_hap1c_eur_sa-qc3* |
| 6 | Cohort 2 (Second QC)<br>sim_hap2a_eur_sa-qc1 | Plink --exclude | hap2a_snp.exc |
| 7 | hap2a_snp.exc | RICOPILI : *preimp* (Default parameters) | sim_hap2b_eur_sa-qc2* |
| 8 | Cohort 3 (Second QC)<br>sim_hap3a_eur_sa-qc1 | Plink --keep | hap3a_only_eur |
| 9 | Cohort 3 (Second QC)<br>hap3a_only_eur | RICOPILI : *preimp* (Default parameters) | sim_hap3b_eur_sa-qc2* |

Extended Data Table 7: Summary of Quality Control Procedures

***Imputation***[42]

In the previous section, we detailed QC steps necessary to prepare genotype datasets for all five cohorts for imputation. Users may either run 'impute' module for each cohort separately in subdirectory /imputation (which is created by default in /qc directory during 'preimp' module) or create symbolic links to each QC'ed files in a new directory and impute all cohorts at the same time.

| **dataset** | **Final QC'd files for imputation** |
|---|---|
| Cohort 1 | sim_hap1c_eur_sa-qc3 |
| Cohort 2 | sim_hap2b_eur_sa-qc2 |
| Cohort 3 | sim_hap3b_eur_sa-qc2 |
| Cohort 4 | sim_hap4a_eur_sa-qc1 |
| Cohort 5 | sim_hap5a_eur_sa-qc1 |

Extended Data Table 8: List of all final post-QC'd genotypes files for all five cohorts for imputation.

For the purpose of the current report, we imputed all cohorts together, although RICOPILI allows users to perform imputation one dataset at a time [43].

We created symbolic links of QC'd files listed in Extended Data Table 8 into an empty directory and started the 'impute' module using below command replacing REFDIR with the file location of a pre-specified directory of the imputation reference panel on the cluster.[44] In the current analysis, we utilized the HRC reference consisting of 54,330 phased haplotypes and 36,678,882 variants.

```
-bash:~> impute_dirsub --refdir REFDIR --minimac3 --minilong
--out hapgen_5cohort
```

By default RICOPILI performs the pre-phasing/imputation stepwise approach using EAGLE/MINIMAC3. As with other modules, we get a success - message after job submission and email on completion. Users are advised to regularly check ~/impute_dir_info for status of the imputation procedures. Upon successful completion of imputation procedures dosage files (with genotype probabilities) and best guess genotypes were available for downstream association analysis.

[42] https://sites.google.com/a/broadinstitute.org/ricopili/imputation
[43] With the script *'my.joinimp2'* (integrated in the RICOPILI distribution) it is possible to merge distinct imputation directories into one (https://docs.google.com/document/d/1xMceim0cMFMMB01Nr9mjHMPtzFAPeahQyEUXAn0xNzQ/).
[44] However if users are on the LISA cluster, and would like to utilize the 1000 genomes reference panel, the directory is as follows /home/gwas/pgc-samples/hapmap_ref/impute2_ref/1000GP_Phase3_sr_0517d

***Reference Building***[45]

We take a moment here to discuss a feature of RICOPILI that is crucial for imputation. In some cases, where users are using the imputation module in RICOPILI for the very first time, the reference panel (mentioned above) needs to be built on the cluster such that pre-phasing, imputation and later, annotations within the postimp module could be carried out properly. The user may build the 1000 genomes reference panel, which is publicly and freely available for use, on the cluster that they are using. The refdir_navi module assist the user for this purpose. Users may use the following command for building the 1000 genomes directory:

```
-bash:~>refdir_navi --vcf_site
ftp://ftp.1000genomes.ebi.ac.uk/vol1/ftp/release/20130502/ --vcf_templ
ALL.chrXXX.phase3_shapeit2_mvncall_integrated_v5a.20130502.genotypes.vcf.gz
--out_templ ALL_v5a.20130502.chrXXX_1KG_0517 --outname 1KG_0517
--sample_root integrated_call_samples_v3.20130502.ALL.panel
```

In certain cases, due to the longer processing time for the reference panel to be set up, users might have to increase the time allocated for the job using --walltimeplus 10 (increase by 10 hours). Also, users who might also encounter initial breaking of the script that renames the .vcf-files and genetic map files downloaded. In such circumstances, users may consider manually downloading the samples using wget and renaming their files with the “loc.” prefix with the mv command. Users may then restart the pipeline using the above commands. We have a separate document describing the reference building in more detail.[46]

***PCA - module***[47]

#### Estimating PCs and relatedness testing

In large-scale genetic studies of thousands of individuals, there is always a possibility of individuals who are genetically related or duplicated within and across cohorts[48]. It is necessary to identify and adjust for (cryptic) relatedness to avoid biased test statistics. In the current section, we demonstrate how RICOPILI would identify 10 individuals duplicated between Cohorts 4 and 5 (Extended Data Table 1), using the ‘pcaer’ module to identify them. After imputation analysis, RICOPILI creates an additional directory/pcaer_sub in the imputation working directory. There, the pipeline would have automatically created symbolic links of imputed best guess genotype versions (“bgs”) for each cohort. Users may start the ‘pcaer’ module using the following command.

```
-bash:~>  pcaer --prefercase --preferfam --out
cobg_gw.hapgen_5cohort_sim_hap1c_eur_sa-qc3.hg19.ch.fl.bgs.bim
```

[45] https://docs.google.com/document/d/1pfJZmacumWHavtWP4aej2685c5gbLVA37rhba6M_vaw/
[46] https://docs.google.com/document/d/1pfJZmacumWHavtWP4aej2685c5gbLVA37rhba6M_vaw/
[47] https://sites.google.com/a/broadinstitute.org/ricopili/pca
[48] You can merge distinct imputation directories to perform a common PCA. See this document here https://docs.google.com/document/d/1xMceim0cMFMMB01Nr9mjHMPtzFAPeahQyEUXAn0xNzQ/

```
sim_hap2b_eur_sa-qc2.hg19.ch.fl.bgs.bim
sim_hap3b_eur_sa-qc2.hg19.ch.fl.bgs.bim
sim_hap4a_eur_sa-qc1.hg19.ch.fl.bgs.bim
sim_hap5a_eur_sa-qc1.hg19.ch.fl.bgs.bim
```

'--prefercase' would retain cases over controls for cross-phenotypic overlap and '--preferfam' would retain trio individuals over case-control individuals

#### File1: cobg_gw.hapgen_5cohort.mepr.overlap.pdf

### 1 caca and coco

#### 1.1 overlap

| ca/co | sim.hap5b |
|---|---|
| sim.hap4b | 2/1 |
| sim.hap5b | - |

| ca/co | sim.hap4b | |
|---|---|---|
| sim.hap4b | - | |
| sim.hap5b | - | - |

#### 1.2 related

| ca/co | sim.hap5b |
|---|---|
| sim.hap4b | - |
| sim.hap5b | - |

| ca/co | sim.hap4b | |
|---|---|---|
| sim.hap4b | - | |
| sim.hap5b | - | - |

### 2 caco and coca

#### 2.1 overlap

| ca/co | sim.hap5b |
|---|---|
| sim.hap4b | 7/0 |
| sim.hap5b | - |

Figure 32: Related individuals between cohorts.

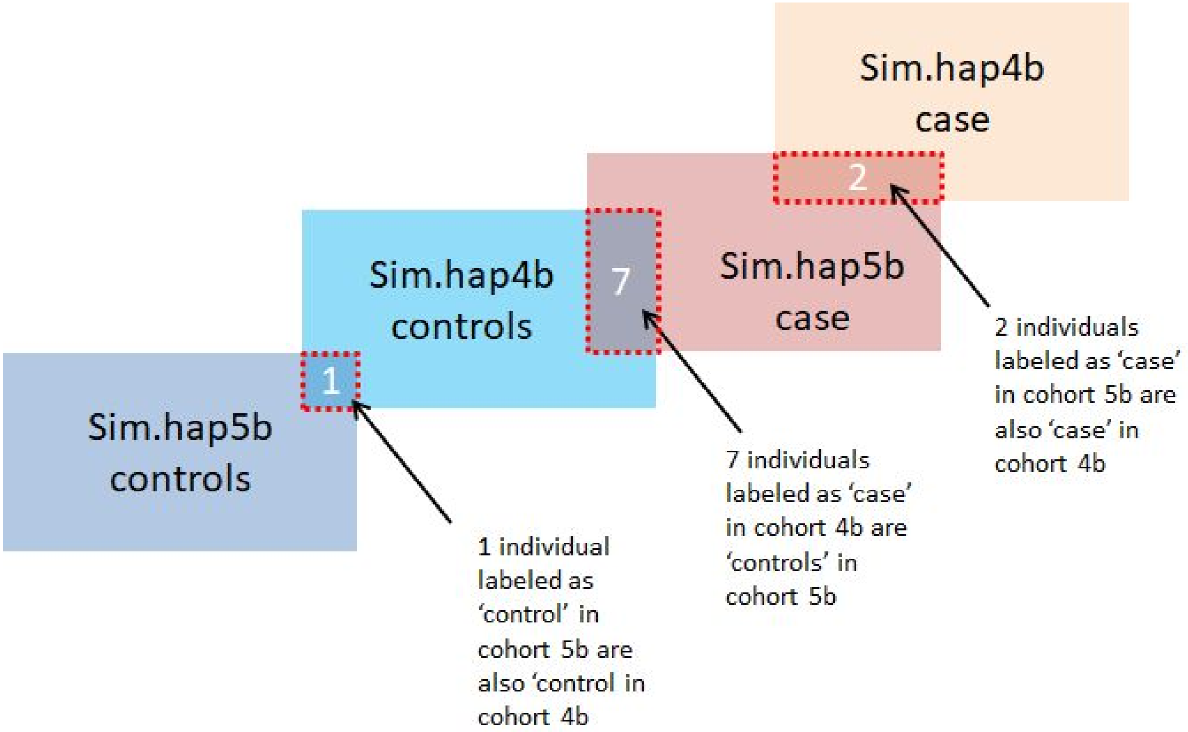


Extended Data Figure 33:Cases and controls overlapping between hap5b and hap4b (interpretation of Figure 32)

In Extended Figures 32 and 33 RICOPILI identified 10 individuals overlapping between hap4b (cohort 4) and hap5b (cohort 5). Two case/case pairs, one control/control pair and 7 discordant pairs are shown in the distinct tables.[49] Essential output files form the PCA module are listed in Extended Data Table 9.

[49] As mentioned above, the flag '--prefercase' excludes the controls for discordant pairs in downstream analyses.

| S.No | Files | Description |
|---|---|---|
| 1 | cobg_gw.hapgen_5cohort.menv.trans.mds | All individuals (non-deduped) with 20 PCAs |
| 2 | cobg_gw.hapgen_5cohort.menv.mds_cov | Deduped subset with 20 PCAs and also study indicator as multiple binary covariates |
| 3 | cobg_gw.hapgen_5cohort.menv.mds | Deduped subset with 20 PCAs without study indicator |
| 4 | cobg_gw.hapgen_5cohort.mepr.overlap | Lists all related / overlapping pairs with IBD0,1,2 and PIHAT |
| 5 | cobg_gw.hapgen_5cohort.mepr.overlap.count | Counts how many an individual has a related/overlapping pair in the collection |
| 6 | cobg_gw.hapgen_5cohort.mepr.overlap.pdf | Tables with study-study numbers (e.g. Figure 32) |
| 7 | cobg_gw.hapgen_5cohort.mepr.famex | Lists the excluded individuals during the deduping process (column 1 and 2) and the individual that has been kept (columns 3 and 4) |
| 8 | cobg_gw.hapgen_5cohort.mepr.genome | Same as *overlap, only with more relaxed PIHAT filter (half of the overlap filter) |
| 9 | cobg_gw.hapgen_5cohort.menv.mds.1d.pdf.gz | Plot of each PC in 1-dimension. |
| 10 | cobg_gw.hapgen_5cohort.menv.mds.2ds.pdf.g z | Plots of first six PC against each other in 2-dimension. |
| 11 | cobg_gw.hapgen_5cohort.menv.mds.2ds.pdf.g z | Shows the PC that are significantly associated with the phenotype (e.g. Figure 34). |

Extended Data Table 9: Important result files of PCA module[50].

[50] https://sites.google.com/a/broadinstitute.org/ricopili/pca#TOC-Output-Files

**File 2: cobg_gw.hapgen_5cohort.menv.mds.sum.pdf**

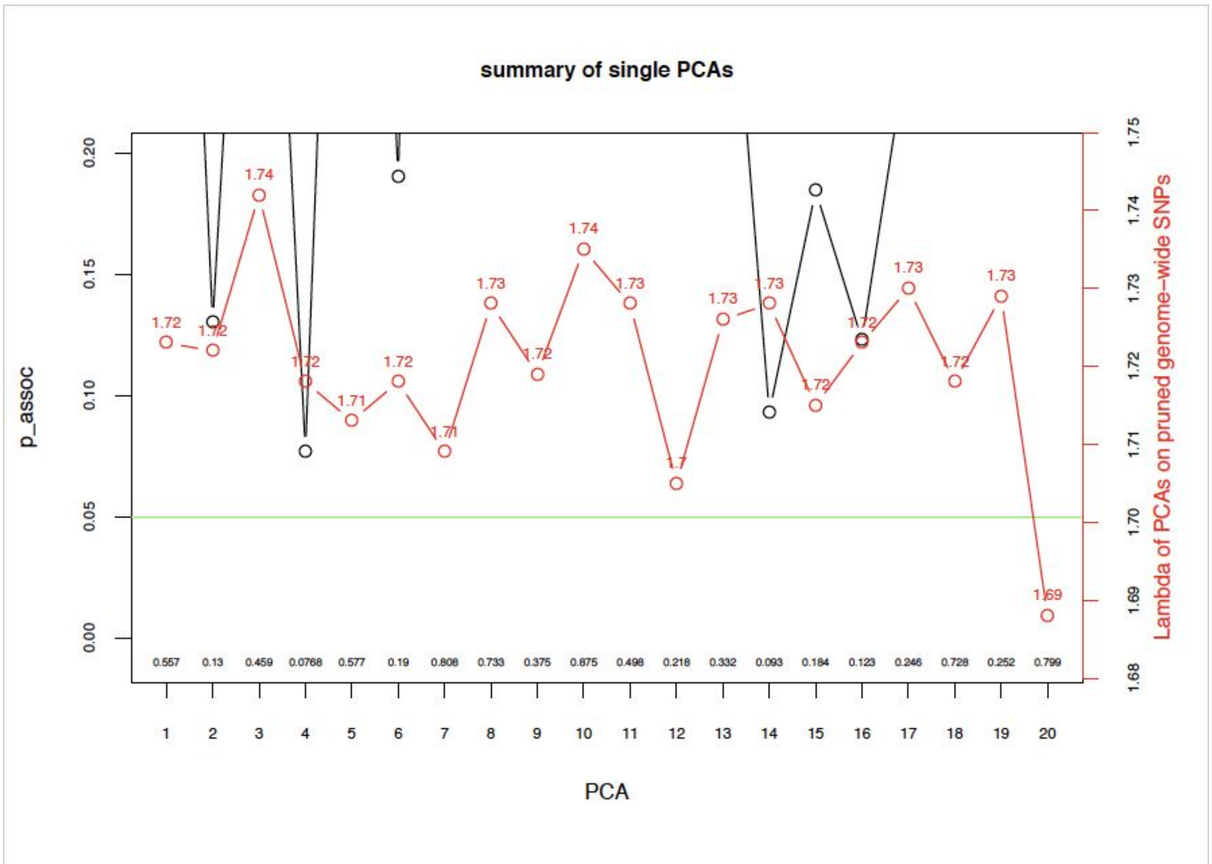


Extended Data Figure 34: List of principal component significantly associated with the phenotype.

RICOPILI conducts association analysis between the first 20 principal components (PCs) with the main phenotype (adjusting for cohort). The PCs were then utilized as covariates in subsequent association testing to adjust for fine-grained population stratification. In Extended Data Figure 34 (black line), we can see that none of the PCs were found to be significantly associated (a real world example could be found here, where PCs 1-7, 9, 13 were significantly associated) with pruned genome-wide SNPs. The red line represents lambda inflation from genome-wide association tests using PCs as phenotype. Depending on the association patterns in this chart, users may stipulate the PCs that they would like to include as covariates in post-imputation analysis using the --coco flag. We recommend including the first four PCs and other associated PCs as covariates.

##### *Post imputation* [51]

#### Genome-wide (meta) association analysis

Use the 'postimp' module to perform genome-wide association analysis of each cohort separately with subsequent meta-analysis and various visualization plots, tables, documents. The 'postimp' module utilized imputed dosage data from the imputation step. Principal components were included in the association analysis as covariates from cobg_gw.hapgen_5cohort.menv.mds_cov

[51] https://sites.google.com/a/broadinstitute.org/ricopili/association-analysis

from previously completed PCA. It is notable that the '_cov' file includes a list of non-duplicated individuals between or within cohorts. In the current analysis PCs 1,2,3,4 were included as covariates. The 'postimp' module was initiated with the command below.

```
-bash:~> postimp_navi --out hapgen_5_cohorts --mds
cobg_gw.hapgen_5cohort.menv.mds_cov --coco 1,2,3,4 --addout
run_2a
```
[52]

In the following sections, we will explain the various output files found in the sub-directory "/distribution/hapgen_5_cohorts_run_11feb/" by RICOPILI. lease also refer to the online references [53]for more details.

##### File: basic.hapgen_5_cohorts_run_2a.num.xls

We provide a file that gives an overview of the sample sizes, distribution of cases/controls, number of SNPs, Genomic inflation (Lambda) and effective sample size for each cohort and included in the combined meta-analysis

| Datasets | N_cases | N_controls | LAMBDA-GC | N-SNPs | N_eff_half |
|---|---|---|---|---|---|
| sim_hap1c_eur_sa-qc3.hg19 | 927 | 965 | 0.991 | 7040031 | 945 |
| sim_hap2b_eur_sa-qc2.hg19 | 423 | 477 | 0.995 | 7078492 | 448 |
| sim_hap3b_eur_sa-qc2.hg19 | 473 | 527 | 0.996 | 7051009 | 498 |
| sim_hap4a_eur_sa-qc1.hg19 | 481 | 514 | 0.993 | 7108825 | 496 |
| sim_hap5a_eur_sa-qc1.hg19 | 516 | 479 | 0.986 | 7107977 | 496 |
| SUM | 2820 | 2962 | 0.982 | 7149025 | 2886 |

Extended Data Table 10: Resulting table form basic file.

[52] --out will simply be added to --out
[53] https://www.dropbox.com/sh/jqp6tc9zph4de6k/AADE-fnNhiJNNbpcgmG0SKwqa?dl=0

**File: qq.hapgen_5_cohorts_run_2a.pdf**

RICOPILI produced a QQ plot showed the p-value distribution of the meta-analysis. Lambda of 0.984 indicate that hidden biases were unlikely.

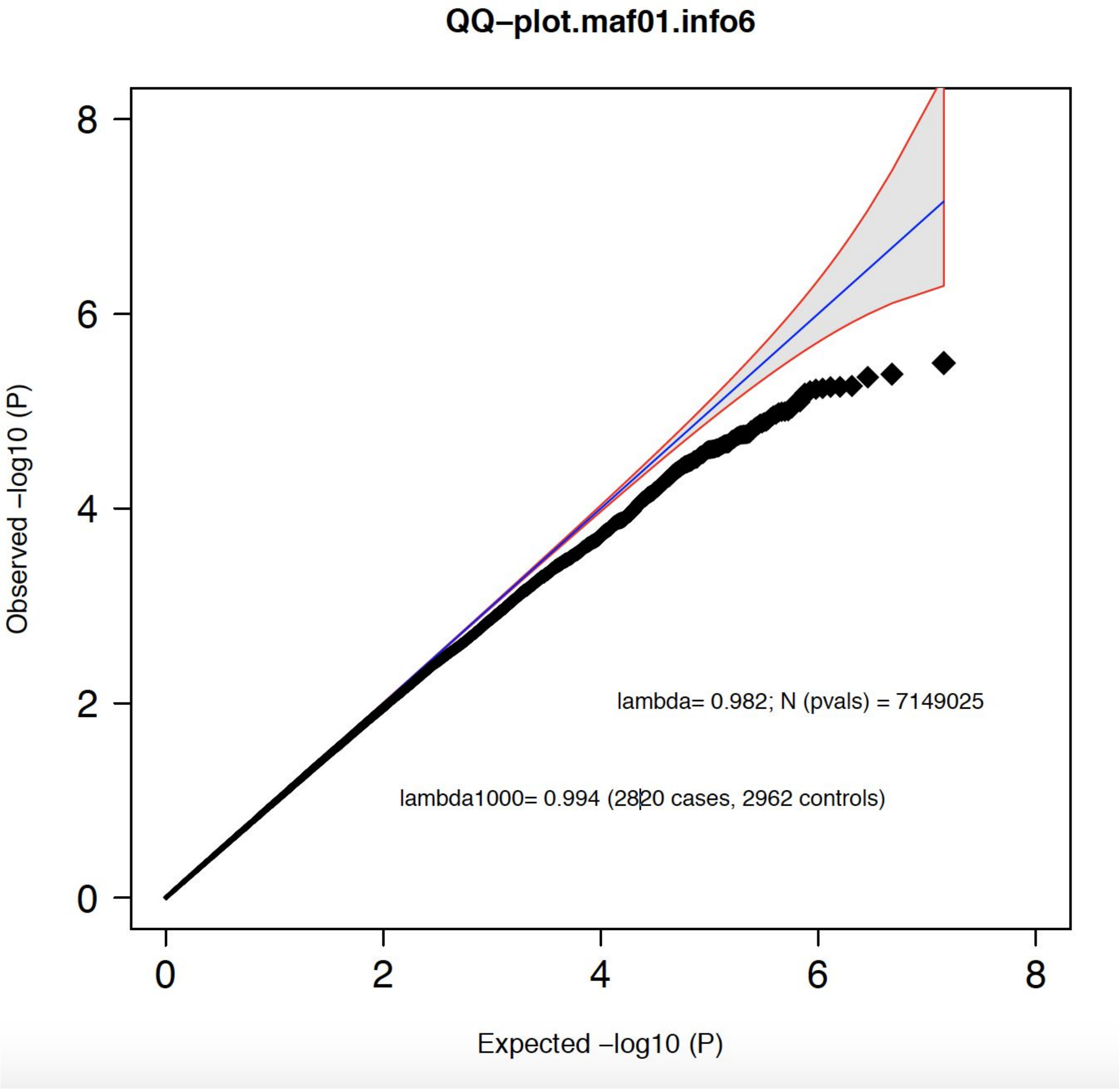


Extended Data Figure 35: QQ plot for overall meta-analysis.

#### File: manhattan.nog2.hapgen_5_cohorts_run_2a.pdf

Various manhattan plots are produced by the pipeline for visualization of the association results after a successful 'postimp' run. An example is provided in Extended Data Figure 36. There are no genome-wide significant SNPs associated with the phenotype.

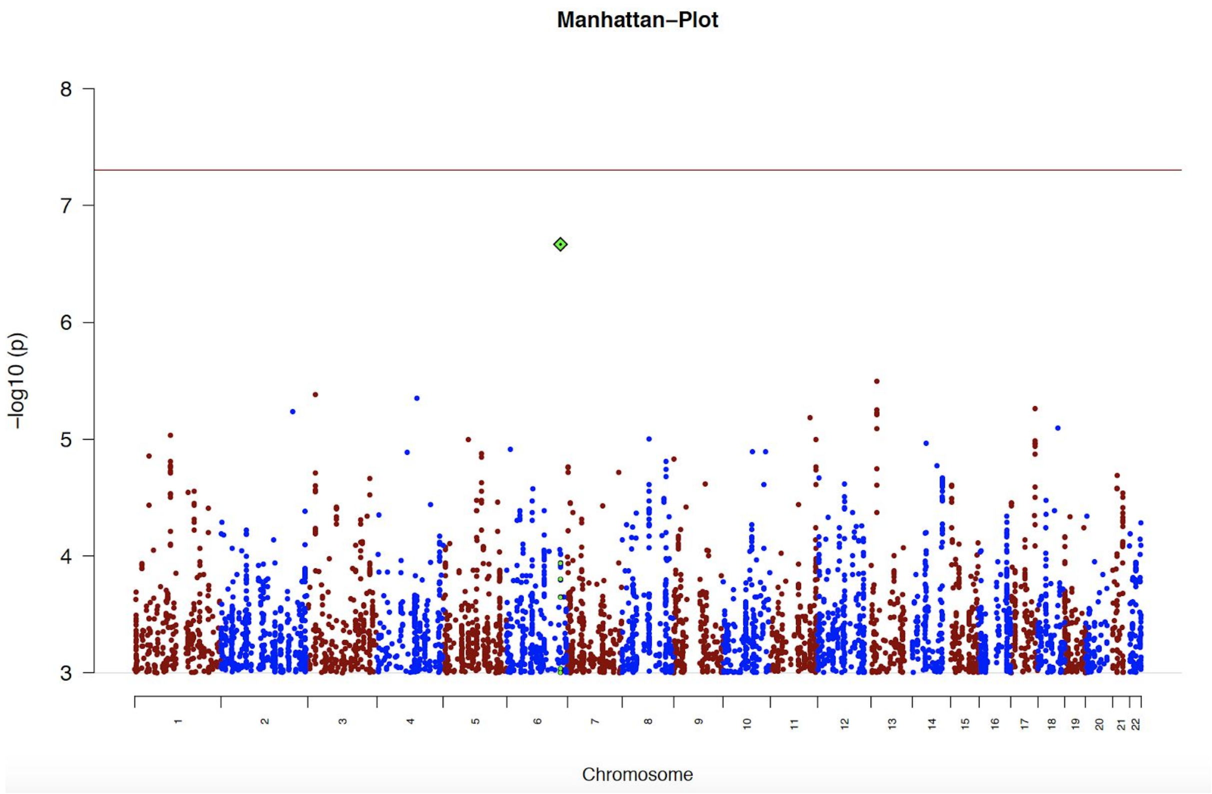


Extended Data Figure 36: Manhattan plot from the meta-analysis.

#### File: areas.hapgen_5_cohorts_run_2a.pdf

Region plots were also produced for genomic regions containing an index SNP with genome-wide significance. As no SNPs met genome-wide significance thresholds, RICOPILI produced the ten top associated regions.

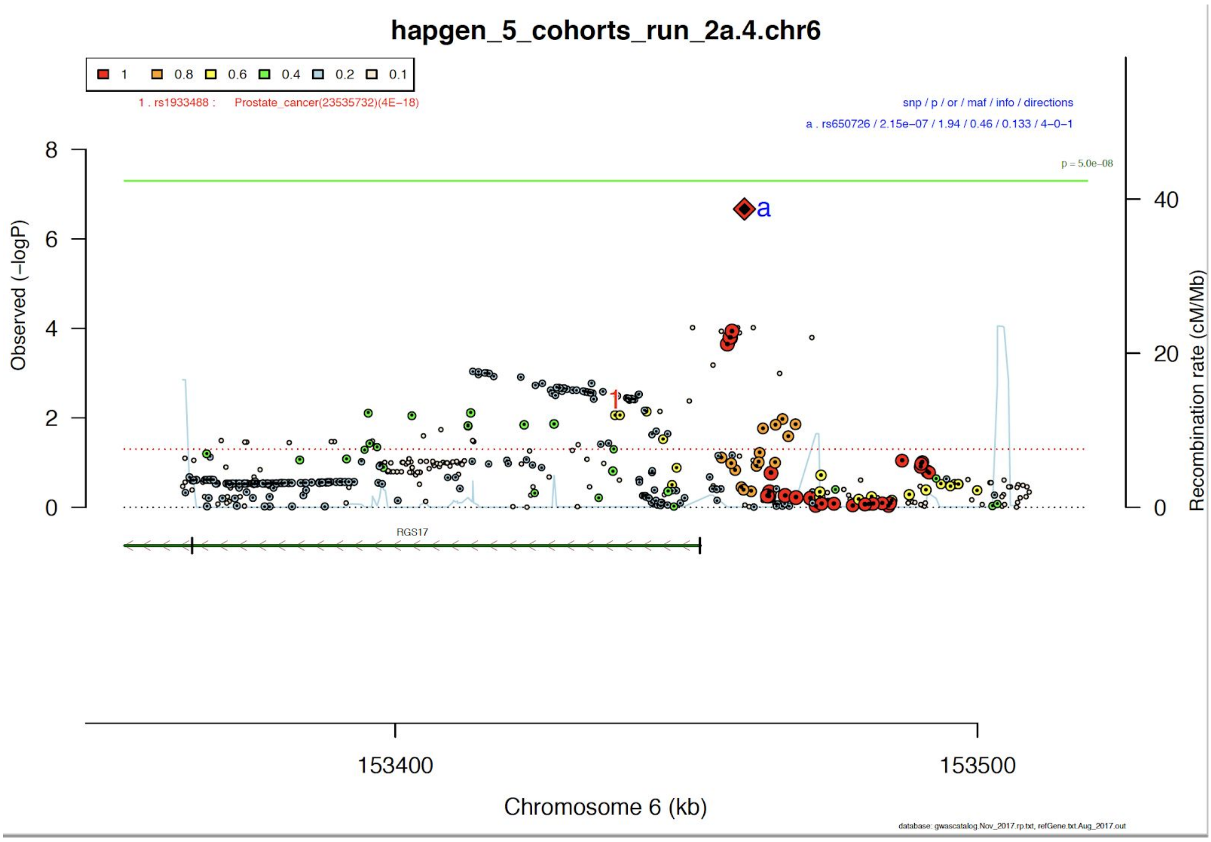


Extended Data Figure 37: Region plot of the r650726, top SNPs from the meta-analysis.

In the current example the index SNP with the lowest p-value rs650726 did not reach genome-wide significance level of $p < 5e-08$. The different colors represent linkage disequilibrium (measured as $r^2$) of each SNP to the index SNP. Most of the SNPs in complete LD (red) with index SNPs show very little evidence for an association, which could point to a false positive, especially in combination with the low imputation quality, INFO score of 0.133.[54] Users are reminded that 'subthreshold' GWAS signals should not be pursued further, without sufficient evidence these could likely be true positives.

---

[54] This region also achieves only a very low region quality score with the RQC method (manuscript in preparation).

#### File: areas.fo.hapgen_5_cohorts_run_2a.pdf

In addition RICOPILI produced forest plots that describe the effect size and its direction in each cohort for each lead SNP. Information such as cohort-specific and meta-analytic info-scores, genotype/imputed SNP, p-value are included in the table.

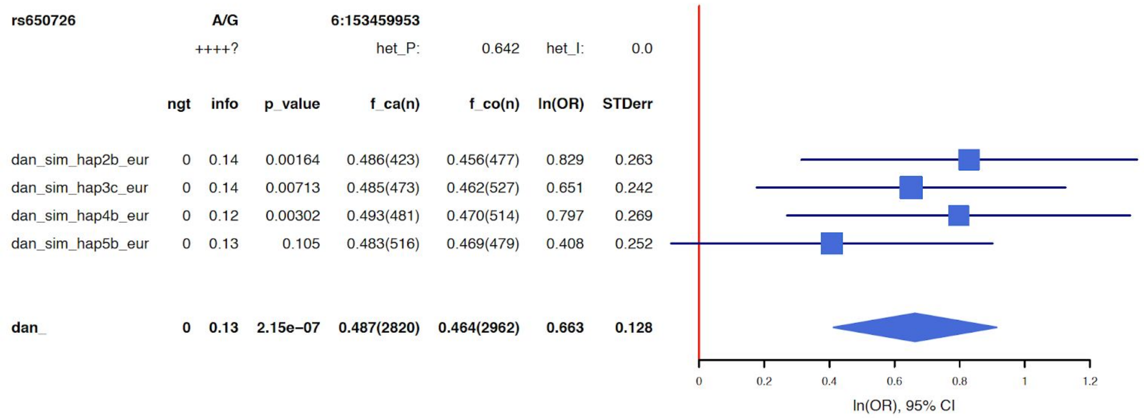


Figure 38: Forest plot of SNP rs650726

Extended Data Figure 38 shows that SNP rs650726 is found in four of the cohorts and missing in one (see directional info: “++++?”). The direction of effect in all the four cohort is same. This SNPs was imputed in all the four cohort (indicated with “0” in column ngt) with a very low info score (INFO < 0.6 is considered low).

#### File: ldsc.daner_hapgen_5_cohorts_run_2a.gz.tar.gz

RICOPILI also performed LD score regression analysis to estimate heritability, intercept and genetic correlation with other phenotypes.

```
Total Observed scale h2: 0.0224 (0.0699)
Lambda GC: 0.9898
Mean Chi^2: 0.9868
Intercept: 0.9842 (0.0071)
```

Extended Data Figure 39: Observed genetic heritability

Summary of Genetic Correlation Results

| p1 | p2 | rg | se | z | p | h2_obs | h2_obs_se | h2_int | h2_int_se | gcov_int | gcov_int_se |
|---|---|---|---|---|---|---|---|---|---|---|---|
| daner_hapgen_5_cohorts_run_2a.gz.ldsc.sumstats.gz | daner_PGC_SCZ49.sh2_mds10.gz.ldsc.sumstats.gz | NA | NA | NA | NA | 0.4707 | 0.0213 | 1.0402 | 0.0132 | 0.0037 | 0.0057 |
| daner_hapgen_5_cohorts_run_2a.gz.ldsc.sumstats.gz | daner_MDD29.0515a_mds6.gz.ldsc.sumstats.gz | NA | NA | NA | NA | 0.1157 | 0.0144 | 0.9836 | 0.0082 | 0.0038 | 0.0046 |

Figure 40: Genetic co-heritability with other phenotype (by default SCZ and MDD)

As expected with the simulated phenotype, SNP heritability was non-significant with observed scale heritability at 2.24% (SE 7%) and there is no co-heritability with schizophrenia and major depression.

##### *Polygenic risk scoring (PRS)*[55]

The 'Postimp' module can also be used for polygenic risk scoring using dosage data. Summary statistics from external sources have to be converted to "daner format" [56] and LD-"clumped" using 'clump_nav3'.

##### *Leave one out (LOO) polygenic risk scores (PRS)*[57]

To perform LOO-PRS we were able to leverage the existing RICOPILI directory structure. The script 'my.preploo' prepares the directory for such analysis. The detailed document can be found here[12].

```
-bash:~>  my.preploo2 --mds
cobg_gw.hapgen_5cohort.menv.mds_cov --postimp postimp_navi
--coco 1,2,3,4 hapgen_5_cohorts_run_2a
```

Once the command is executed, the 'postimp' procedure will guide the user through setup and execution of the LOO analysis (as many as there are cohorts in your collection). It excludes each cohort, performs meta-analysis on remainder, clumps the results and scores the individuals in the excluded cohort. Here, we carried out LOO analysis on each of the cohorts that were imputed. Results of the LOO analysis is reported in Extended Data Figure 41.

[55] https://docs.google.com/document/d/10jvLvnrPIz9zRKlRfMeW8HsHDxEIbs84yP3Ugjhj31E/
[56] https://docs.google.com/document/d/1TWIhr8-qpCXB13WCXcU1_HDio8lC_MeWoAg2jlggrtU/
[57] https://docs.google.com/document/d/10UKUfX4wT9rP5zswH5SDR2Y1O2-wJVUw9mn4tSdaS-I/

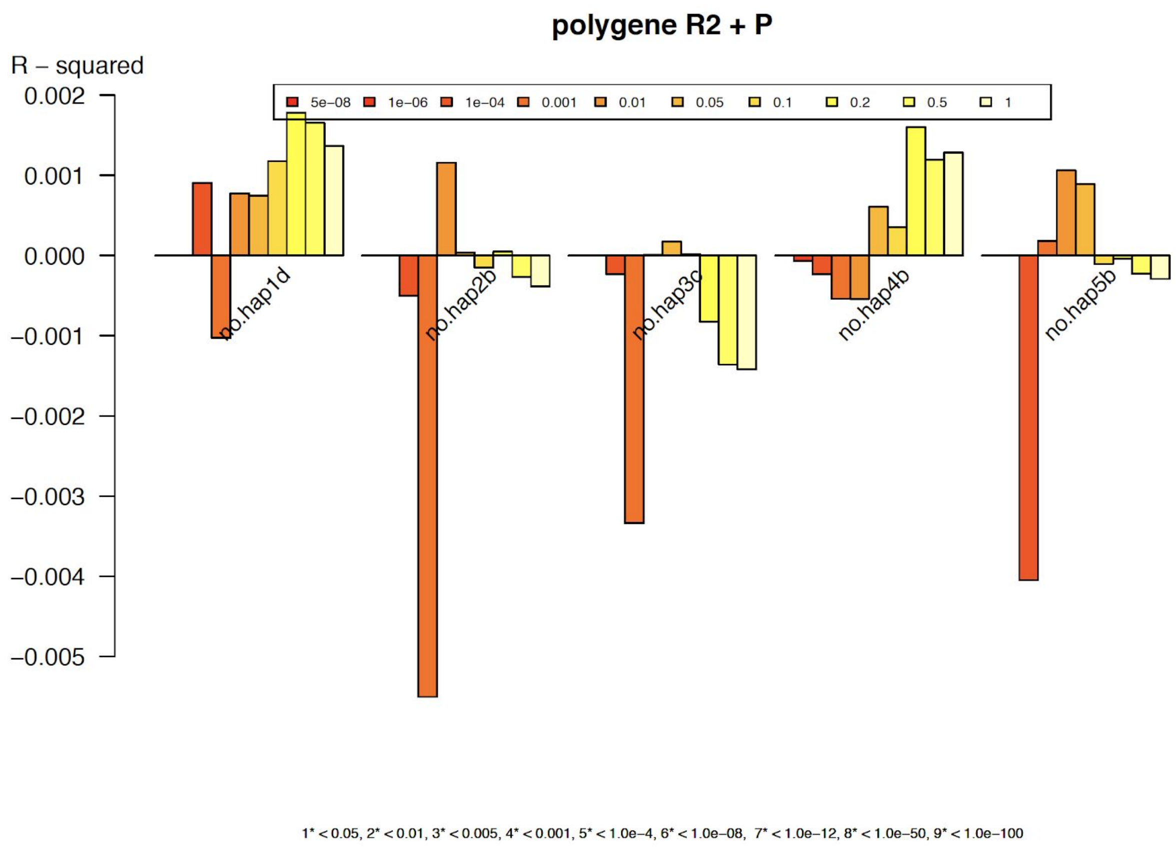


Extended Data Figure 41: LOO analysis results; y-axis is variation explained (measured as nagelkerke r2) at various p-value cut-of in each cohort on x-axis.

Association (measured as nagelkerke r2) between risk scores and phenotype were not significant and a random direction of effect was observed across the simulated phenotypes.

#### *Replication*

The RICOPILI output files and directory structure was also designed to facilitate replication analysis with ease[58]. To demonstrate the functionality of the replication module, we carried out LOO analysis as reported in the previous section[59]. Specifically, replication analysis was carried out for cohort hapgen4b, while using the meta-analysis results for the other three cohorts. The following three files produced by RICOPILI were crucial for the replication analysis:

a. daner_hapgen_5_cohorts_run_11feb.loo.no.hap4b.gz.p4.clump.areator.sorted.1mhc ( This contains associated genomic regions at P<1e-4. That have been clumped to account for linkage disequilibrium amongst SNPs).
b. daner_hapgen_5_cohorts_run_11feb.loo.no.hap4b.gz.p3.gz (Meta-analysis results for all SNPs with P < 1e-3).
c. daner_sim_hap4b_eur_sa-qc.hg19.ch.fl.gz.p3.gz. (Cohort 4 association results).

[58] https://docs.google.com/document/d/1mvr2Kx2MrYaqlVIlE5nSpMePTO_qWkmSZySdT4AdBQw/
[59] We can use meta-analysis from different sources convert them into daner format or generate required file using *'posimp'* module with --*result* flag. If the replication data is not in daner format (use --format 0) in the *'replicator13'* module.

Files 'a' and 'b' could be found in the following directories:
results_hapgen_5_cohorts_run_2a/distribution/hapgen_5_cohorts_run_2a.loo.no.hap4b
sub-directory from LOO-PRS step

and the file 'c' can be found in /report_hapgen_5_cohorts_run_2a from the 'postimp' directory.

We copied these files in new directory and started 'replicator13' module in an interactive shell using the command below. (Note: Replace REFDIR with imputation reference directory),

```
-bash:~> replicator13 --dan --areator
daner_hapgen_5_cohorts_run_2a.loo.no.hap4b.gz.p4.clump.areator
.sorted.1mhc --gwas
daner_hapgen_5_cohorts_run_2a.loo.no.hap4b.gz.p3.gz --format 7
--refdir REFDIR --index --trust --rep
daner_sim_hap4b_eur_sa-qc.hg19.ch.fl.gz
```

We explain briefly the functions some of the flags we utilized for the replication analysis: --index searches for index SNP (not any LD-friends) in the --rep i.e replication data. --trust indicates that SNPs were assumed to be on the same strand and frequency checks were omitted on ambiguous SNPs. The results of the replication analysis results were combined into a single excel file; An Example File is here for download.[60] For detailed description of the replication analysis results file format please refer here. Sign test results are described in Extended Data Table 11. There were no significant sign test results.

| Name | P_TH | npos | nsum | sign-test | ratio | Ncase | Ncontrol |
|---|---|---|---|---|---|---|---|
| hapgen_5_cohorts_run_2a.loo.no.hap4b.sim_hap4b_eur_sa-qc.hg19.ch.fl.rep | 0.0001 | 60 | 127 | 0.761039 | 0.47 | 481 | 514 |
| hapgen_5_cohorts_run_2a.loo.no.hap4b.sim_hap4b_eur_sa-qc.hg19.ch.fl.rep | 0.00001 | 5 | 12 | 0.806152 | 0.42 | 481 | 514 |
| hapgen_5_cohorts_run_2a.loo.no.hap4b.sim_hap4b_eur_sa-qc.hg19.ch.fl.rep | 0.000001 | 0 | 1 | 1 | 0 | 481 | 514 |
| hapgen_5_cohorts_run_2a.loo.no.hap4b.sim_hap4b_eur_sa-qc.hg19.ch.fl.rep | 5E-08 | 0 | 0 | 1 | NA | 481 | 514 |
| REPLICATION | 0.0001 | 60 | 127 | 0.761039 | 0.47 | 481 | 514 |
| REPLICATION | 0.00001 | 5 | 12 | 0.806152 | 0.42 | 481 | 514 |
| REPLICATION | 0.000001 | 0 | 1 | 1 | 0 | 481 | 514 |
| REPLICATION | 5E-08 | 0 | 0 | 1 | NA | 481 | 514 |
| REPLICATION | 1E-08 | 0 | 0 | 1 | NA | 481 | 514 |
| REPLICATION | 5E-09 | 0 | 0 | 1 | NA | 481 | 514 |
| REPLICATION | 5.0e-08_to | 0 | 0 | 1 | NA | 481 | 514 |
| REPLICATION | 1.0e-06_to | 0 | 1 | 1 | 0 | 481 | 514 |
| REPLICATION | 1.0e-05_to | 5 | 11 | 0.725586 | 0.45 | 481 | 514 |

Extended Data Table 11: Replication/Sign test results

[60] https://docs.google.com/spreadsheets/d/1ISvOMHM7IwXhbTXMqb4NUvDMITKqEqRCeC2uLvi--k4/
